# Thalamofrontal synaptic weakening underlies short-term memory deficits from adolescent NMDAR hypofunction

**DOI:** 10.1101/2025.11.11.687794

**Authors:** Jinseon Yu, In Sun Choi, Gyu Hyun Kim, Sangkyu Bahn, Sungwon Bae, Taekwan Lee, Joon Ho Choi, Jun Soo Kwon, Minah Kim, Kea Joo Lee, Jong-Cheol Rah

## Abstract

N-methyl-D-aspartate receptor (NMDAR) hypofunction is linked to schizophrenia, but prevailing models centered on reduced number and activity of inhibitory interneurons, and the resulting excitation–inhibition (E-I) imbalance, do not explain selective cognitive impairments. Here, we show that repeated adolescent NMDAR antagonism produces short-term memory (STM) deficits by weakening thalamofrontal (TF) synaptic transmission. In mice repeatedly exposed to ketamine, STM impairment coincided with reduced release probability and vesicle refilling at mediodorsal thalamus (MD) → dorsomedial prefrontal cortex (dmPFC) synapses, without detectable changes in layer 2/3 → layer 2/3 synaptic release probability, intrinsic excitability, or synaptic ultrastructure. These presynaptic deficits were accompanied by diminished direction-selective population coding in the dmPFC, as revealed by decoding analysis, and impaired delayed alternation performance in the Y-maze. Chemogenetic strengthening of MD → dmPFC projections restored both neural selectivity and behavior. Our findings identify a circuit-specific presynaptic mechanism linking adolescent NMDAR hypofunction to cognitive dysfunction, challenging interneuron-centric models and establishing TF synapses as a potential therapeutic target in NMDAR-related disorders.

## Introduction

N-methyl-D-aspartate receptor (NMDAR) hypofunction has been consistently observed in several neuropsychiatric disorders, including schizophrenia^1–5^. In particular, accumulating clinical evidence indicates that persistent NMDAR dysfunction is one of the core pathophysiological features of schizophrenia^6^. Analyses of cerebrospinal fluid and postmortem brain tissue from patients with schizophrenia have revealed multiple molecular abnormalities consistent with NMDAR hypofunction, including reduced levels of D-serine, a co-agonist essential for NMDAR activation, and decreased mRNA and protein expression of GluN1, the obligatory subunit of the receptor^7–9^. Additionally, trafficking of NR2B-containing NMDARs appears impaired in the prefrontal cortex (PFC)^10,11^, and kynurenic acid—an endogenous NMDAR antagonist—is markedly elevated in schizophrenia patient cerebrospinal fluid compared to healthy controls^12^. These converging findings support the notion that sustained reductions in NMDAR signaling contribute to cognitive and synaptic dysfunction in schizophrenia. To further support this link, administration of NMDAR antagonists such as phencyclidine, ketamine, and MK-801 has been shown to transiently evoke schizophrenia-like symptoms, including short-term memory (STM) impairments, in healthy individuals^13–15^, and to exacerbate symptoms in patients^6,16,17^. However, these pharmacological effects are short-lived and do not capture the long-term circuit-level alterations characteristic of the disorder. To address this gap, we examined the consequences of sustained NMDAR hypofunction in the dorsomedial prefrontal cortex (dmPFC) during adolescence.

Our rationale was threefold: (1) NMDARs are critical for activity-dependent pruning, spine stabilization, and synaptic integration during developmental critical periods^18,19^, (2) the maturation of prefrontal cortex is protracted relative to other cortical regions, extending into adolescence and early adulthood^20–25^. (3) schizophrenia most commonly emerges during adolescence, when prefrontal circuits are still maturing^26,27^. On this basis, we hypothesized that persistent NMDAR antagonism during adolescence—a window of heightened circuit plasticity—can induce enduring changes in cortical connectivity that can account for STM deficits.

To support our hypothesis, animal models of chronic NMDAR hypofunction recapitulate many behavioral and cognitive phenotypes observed in schizophrenia, such as STM deficits, attentional impairments, and disrupted social interaction^28–35^. However, it remains largely unclear how sustained NMDAR hypofunction alters prefrontal circuit function to produce specific cognitive impairments such as STM dysfunction.

We focused on STM deficits because they are among the most consistently impaired cognitive domains in schizophrenia^36^. The severity of STM impairment also correlates with broader symptomatology in schizophrenia patients^37,38^. Furthermore, STM critically depends on fine circuit wiring that supports persistent neural activity within the PFC^39–44^.

While intrinsic mechanisms such as muscarinic receptor activation may contribute to maintaining persistent activity during STM^45^, converging evidence emphasizes the importance of long-range circuits, particularly reciprocal MD–PFC loops, in supporting STM^42,46–48^. Disruption of MD inputs destabilizes prefrontal representations and impairs STM^46,49^. Consistent with this, deficits in thalamofrontal (TF) connectivity have been proposed as a core feature of “cognitive dysmetria” in schizophrenia^50^ and are supported by imaging and electrophysiological studies in patients^51–58^. Complementary loss-of-function experiments across species further demonstrate that interfering with MD activity impairs PFC coding and STM performances^46,49,59,60^.

Here, we tested this hypothesis using a mouse model of repeated ketamine administration. We specifically asked whether sustained NMDAR hypofunction during adolescence disrupts TF (MD → dmPFC) synaptic function to produce STM impairments. Our findings reveal a circuit-specific presynaptic mechanism underlying STM dysfunction and point to TF connectivity as a potential therapeutic target for the cognitive symptoms of schizophrenia and related disorders.

## Results

### STM performance under chronic NMDAR hypofunction

To model adolescent NMDAR hypofunction, we administered intraperitoneal ketamine (30 mg/kg) daily for 14 consecutive days beginning at approximately postnatal day 21 (Fig. 1A). STM performance was assessed using a reward-motivated delayed Y-maze task. In each trial, mice were guided into one of the two randomly selected arms of the maze (sample phase) and confined by a motorized door during a 30-s delay (delay phase). After the delay, the door was opened, and mice were required to enter the opposite, previously unvisited arm to obtain a food pellet (correct trial). Returning to the same arm was defined as an error and was not rewarded. To avoid confounding acute effects, behavioral testing was conducted 24 hours after the final injection, well beyond serum half-life of ketamine^61^. Mice treated with repetitive ketamine administration (rKet) exhibited significantly lower correct choice rates across trials compared to vehicle-treated controls (Ctrl) (Fig. 1B).

**Figure 1.**
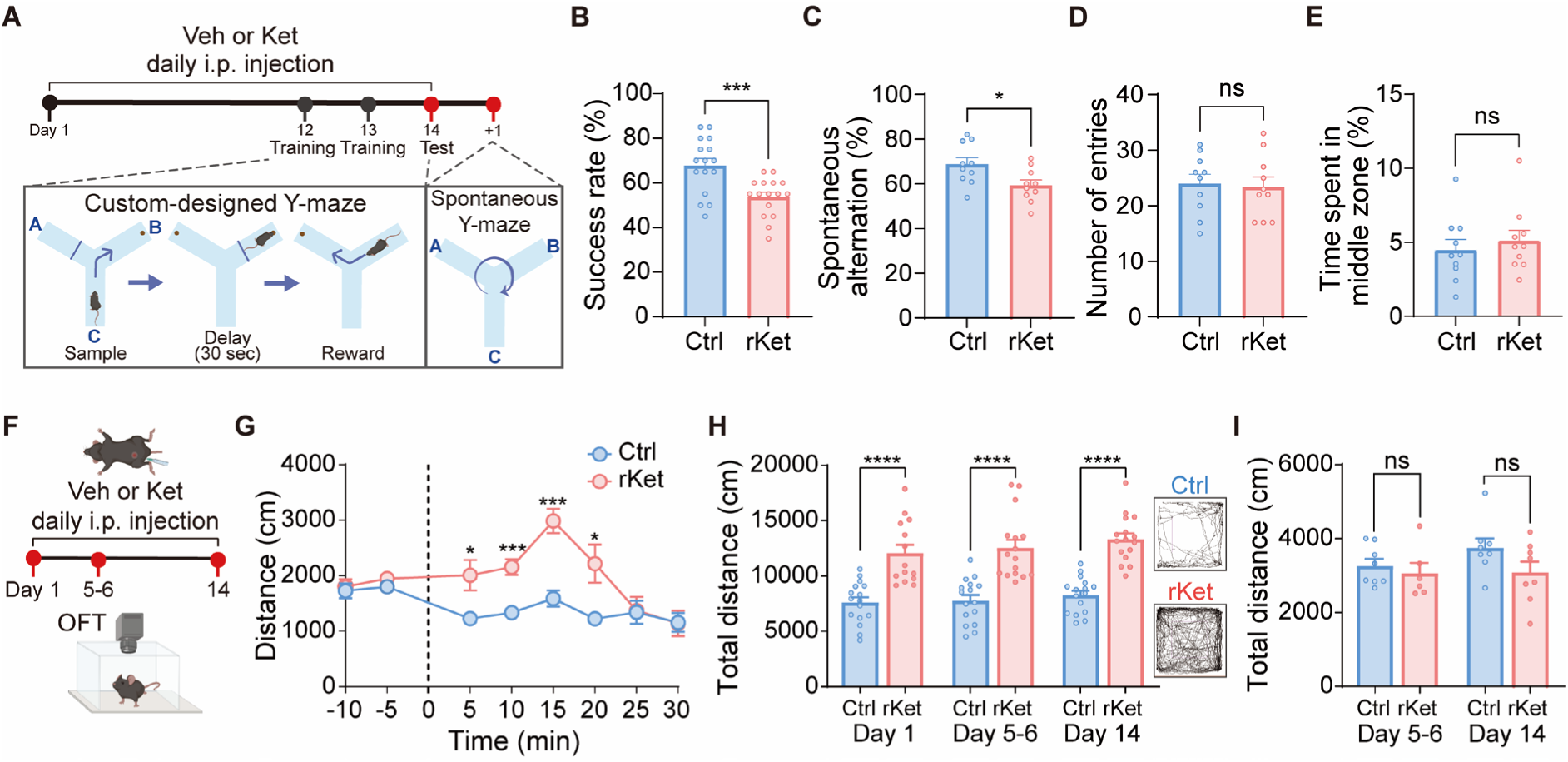
STM performance following chronic NMDAR hypofunction. **A–E,** Experimental design and behavioral assessment of short-term memory (STM). **A,** Experimental schedule and behavioral task design. **B,** Success rate in the custom-designed Y-maze task. The training day consisted of two sessions (morning and afternoon), each comprising 10 trials. The success rate represents the average percentage of correct choices across the two sessions on the test day (Ctrl: repeatedly treated with vehicle, 16 mice; rKet: repeatedly treated with ketamine, 16 mice). **C–E,** Spontaneous alternation performance in the Y-maze (Ctrl: 10 mice, rKet: 10 mice). **C,** Spontaneous alternation performance. **D,** Total number of arm entries. **E,** Percentage of time spent in the middle zone. **F–I,** Assessment of locomotor activity following drug administration using the open field test (OFT). **F,** Experimental scheme for OFT. **G,** The time course of locomotor activity over 30 min following drug administration in OFT. Distance traveled was measured in 5-min bins on Day 1 (Ctrl: 8 mice, rKet: 8 mice). **H,** Acute effects of ketamine immediately after administration in the open field. Left: Total distance during 30 min following drug administration. Right: Representative trajectories from Ctrl and rKet mice at peak hyperlocomotion (Ctrl: 16 mice, rKet: 14 mice). **I,** Summed locomotion distance with repeated ketamine exposure measured 24 hours after the final injection in the open field. Total distance during 10 min before drug administration on day 5-6 and day 14 (Ctrl: 8 mice, rKet: 6 mice). Data are presented as mean ± SEMs; \**p* < 0.05, \*\*\**p* < 0.001, \*\*\*\**p* < 0.0001 and ns, not significant by unpaired Student’s t-test compared with the Ctrl group.

Although this task was designed to assess the effects of rKet on STM, it was important to control potential confounding factors. First, reduced success rate could reflect slower learning, as the modified Y-maze required two sessions of training to reach stable performance (Extended Data Fig. 1) and impaired learning has been reported in schizophrenia patients^62,63^. Second, rKet might induce psychomotor slowing similar to that observed in patients^64,65^, which could effectively increase the memory delay. Third, motivational deficits due to altered effort–value computations^66,67^ could produce similar behavioral outcomes. To address these possibilities, we employed a spontaneous alternation version of the Y-maze that does not require training or reinforcement. rKet mice again showed significantly reduced alternation performance (Fig. 1C), while the number of arm entries per session was comparable between groups (Fig. 1D). The proportion of time spent in the center zone, corresponding to the delay interval, was also unchanged (Fig. 1E). Together, these results indicate that the reduced performance reflects an STM deficit rather than reduced locomotor activity, exploratory drive, or engagement.

We next tested whether the behavioral changes reflected chronic rather than acute effects of ketamine (Fig. 1F). As expected, ketamine induced a transient hyperlocomotor response that peaked within 20 minutes post-injection (Fig. 1G–H), consistent with its known pharmacokinetics in rodents (serum half-life ∼13–20 minutes^61^). However, locomotor or anxiety activity assessed 24 hours after injection—the time point used for behavioral task—was indistinguishable from controls (Fig. 1I, Extended Data Fig. 2), indicating that reduced STM performance results from long-term effects of repeated ketamine exposure rather than acute pharmacological action. Furthermore, STM performance was significantly reduced at 1 day after the final rKet injection and did not show recovery even after 28 days (Extended Data Fig. 3), indicating a persistent impairment.

### Spontaneous excitatory and inhibitory synaptic transmission to layer 2/3 pyramidal neurons in the dmPFC

We next examined whether repeated NMDAR hypofunction alters synaptic transmission in the dmPFC, a region critical for STM. Previous studies have suggested that NMDAR antagonism preferentially suppresses the activity of fast-spiking interneurons, leading to disinhibition and elevated pyramidal firing rates^68^. Consistent with previous reports in both rodents and patients with schizophrenia^31–33,69,70^, rKet treatment led to a significant reduction in the density of parvalbumin-positive (PV^+^) interneurons in dmPFC, visualized using PV-tdTomato transgenic mice (Fig. 2A–B). Accordingly, whole-cell recordings from layer 2/3 pyramidal neurons revealed a decrease in the frequency, but not amplitude, of spontaneous inhibitory postsynaptic currents (sIPSCs), consistent with reduced inhibitory input without postsynaptic receptor changes (Fig. 2C). Unexpectedly, a significant decrease in the frequency of spontaneous excitatory postsynaptic currents (sEPSCs) was observed in rKet mice (Fig. 2D).

**Figure 2.**
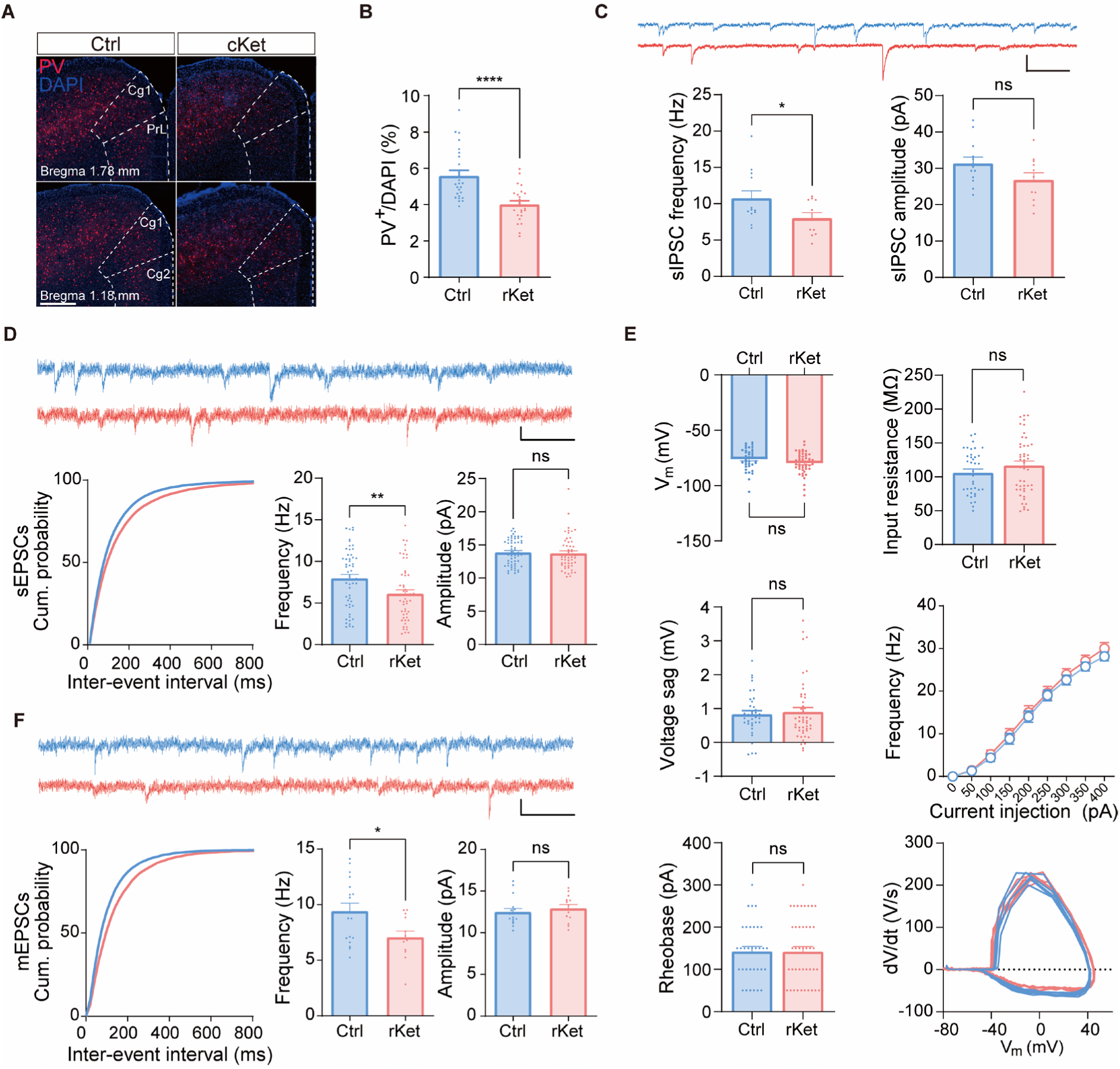
Spontaneous synaptic transmission and intrinsic properties of layer 2/3 pyramidal neurons in the dmPFC. **A,** Representative fluorescent images showing PV^+^ interneurons (red) and DAPI (blue) in the dorsomedial prefrontal cortex (dmPFC) of PV-tdTomato mice following drug administration. Scale bar: 500 μm. **B,** Quantification of PV^+^/DAPI neuron percentage in dmPFC. We analyzed 24 slices in total (6 slices per mouse, 4 mice per group). **C,** Quantification of spontaneous inhibitory postsynaptic currents (sIPSCs) recorded from layer 2/3 neurons in the dmPFC following repeated drug administration. Upper: Representative sIPSC traces from Ctrl (blue) and rKet (red) groups. Scale bars: 100 ms, 10 pA. Lower: Average frequency and amplitude from dmPFC layer 2/3 pyramidal neurons (Ctrl: 13 cells from 4 mice; rKet: 11 cells from 4 mice). **D,** Quantification of spontaneous excitatory postsynaptic currents (sEPSCs) recorded from layer 2/3 neurons in the dmPFC following repeated drug administration (Ctrl: 56 cells from 17 mice; rKet: 53 cells from 16 mice). Upper: Representative sEPSC traces from Ctrl (blue) and rKet (red) groups. Scale bars: 100 ms, 10 pA. Lower: Cumulative probability plots of inter-event intervals and summary bar graphs of mean frequency and amplitude. **E,** Intrinsic excitability properties of layer 2/3 pyramidal neurons in the dmPFC: Resting membrane potential (V_m_), input resistance, and voltage sag (Ctrl: 37 cells from 11 mice; rKet: 48 cells from 11 mice), as well as action potential frequencies in response to current injection (0–400 pA in 50 pA steps for 600 ms) and rheobase (Ctrl: 33 cells; rKet: 44 cells from the same cohort of mice). Phase-plane plots (dV/dt vs. membrane potential) show representative cells during current injections at rheobase (150 pA). **F,** Quantification of miniature excitatory postsynaptic currents (mEPSCs) recorded in the presence of tetrodotoxin (TTX, 0.5 μM), from layer 2/3 pyramidal neurons in the dmPFC following repeated drug administration (Ctrl: 17 cells from 6 mice; rKet: 13 cells from 4 mice). Upper: Representative mEPSC traces from Ctrl (blue) and rKet (red) groups. Scale bars: 100 ms, 10 pA. Lower: Cumulative probability plots of inter-event intervals and summary bar graphs of mean frequency and amplitude. Data are presented as mean ± SEMs; \**p* < 0.05, \*\**p* < 0.01, \*\*\*\**p* < 0.0001, and ns, not significant by unpaired Student’s t-test compared with the Ctrl group. Statistical significance for cumulative probability plots was determined by Kolmogorov-Smirnov test \*\*\*\**p* < 0.0001 for panel D and E.

To determine whether intrinsic excitability contributed to these changes, we measured passive and active membrane properties of layer 2/3 pyramidal neurons in the dmPFC. Resting membrane potential, input resistance, voltage sag, firing rates in response to current injection, and rheobase were all unaltered in rKet mice (Fig. 2E), indicating that intrinsic excitability remained intact.

Instead, the reduction in excitatory synaptic input appeared to be presynaptic in origin. Miniature EPSCs (mEPSCs), recorded in the presence of tetrodotoxin (TTX) to block action potentials, were also significantly reduced in frequency (Fig. 2F). Amplitudes of both sEPSCs and mEPSCs were unchanged, arguing against postsynaptic receptor downregulation. Together, these results indicate that repeated NMDAR hypofunction impairs excitatory synaptic transmission via a presynaptic mechanism, without altering intrinsic excitability or E–I balance in the dmPFC.

The reduction in excitatory transmission was a long-lasting effect of ketamine exposure The reduction in excitatory transmission appears to develop gradually with repeated ketamine exposure. After 5 days of administration, cumulative probability analysis revealed a subtle but significant shift in inter-event intervals, although the mean sEPSC frequency was not significantly altered (Extended Data Fig. 4). These results suggest that the decrease in excitatory drive may emerge progressively over time. Interestingly, these effects were region-specific, as neither excitatory synaptic transmission nor intrinsic excitability was altered in the primary somatosensory cortex (S1BF) of rKet mice (Extended data Fig. 5), suggesting that repeated NMDAR hypofunction preferentially affects the dmPFC.

### Release probability and vesicle organization in dmPFC excitatory synapses

To directly assess presynaptic function in the dmPFC under chronic NMDAR hypofunction, we performed whole-cell recordings from layer 2/3 pyramidal neurons and examined short-term plasticity using perisomatic stimulation of local inputs. To minimize potential postsynaptic contributions to short-term plasticity, such as receptor saturation^71,72^, we adjusted stimulation intensity to evoke comparable initial EPSC amplitudes across groups (Ctrl: 185.9 ± 11.7 pA; rKet: 182.8 ± 12.3 pA), ensuring equalized postsynaptic involvement (Fig. 3C). At 10 Hz stimulation, control neurons displayed prominent short-term depression, whereas this depression was significantly attenuated in rKet mice (Fig. 3D–E), suggesting a reduction in release probability (Pr). To further dissect the mechanism underlying this reduced release efficacy, we applied a high-frequency (33.3 Hz) stimulus train to deplete the readily releasable pool (RRP) and constructed cumulative EPSC amplitude plots. Using an established approach by Schneggenburger and colleagues, the slope of the late linear phase was used to estimate the vesicle refilling rate, while the y-intercept of the extrapolated fit provided an estimate of RRP size (Fig. 3F). Pr was then calculated as the ratio of the first EPSC amplitude to the estimated RRP. Both refilling rate and Pr were significantly reduced in rKet mice compared to controls (Fig. 3G-H), indicating that chronic NMDAR hypofunction impairs both the initial probability of vesicle release and the replenishment of release-ready vesicles. Although the estimated RRP size appeared similar between groups (Fig. 3I), this interpretation should be taken with caution, as stimulation intensity was normalized to evoke similar initial EPSCs, potentially masking group differences in absolute pool size.

**Figure 3.**
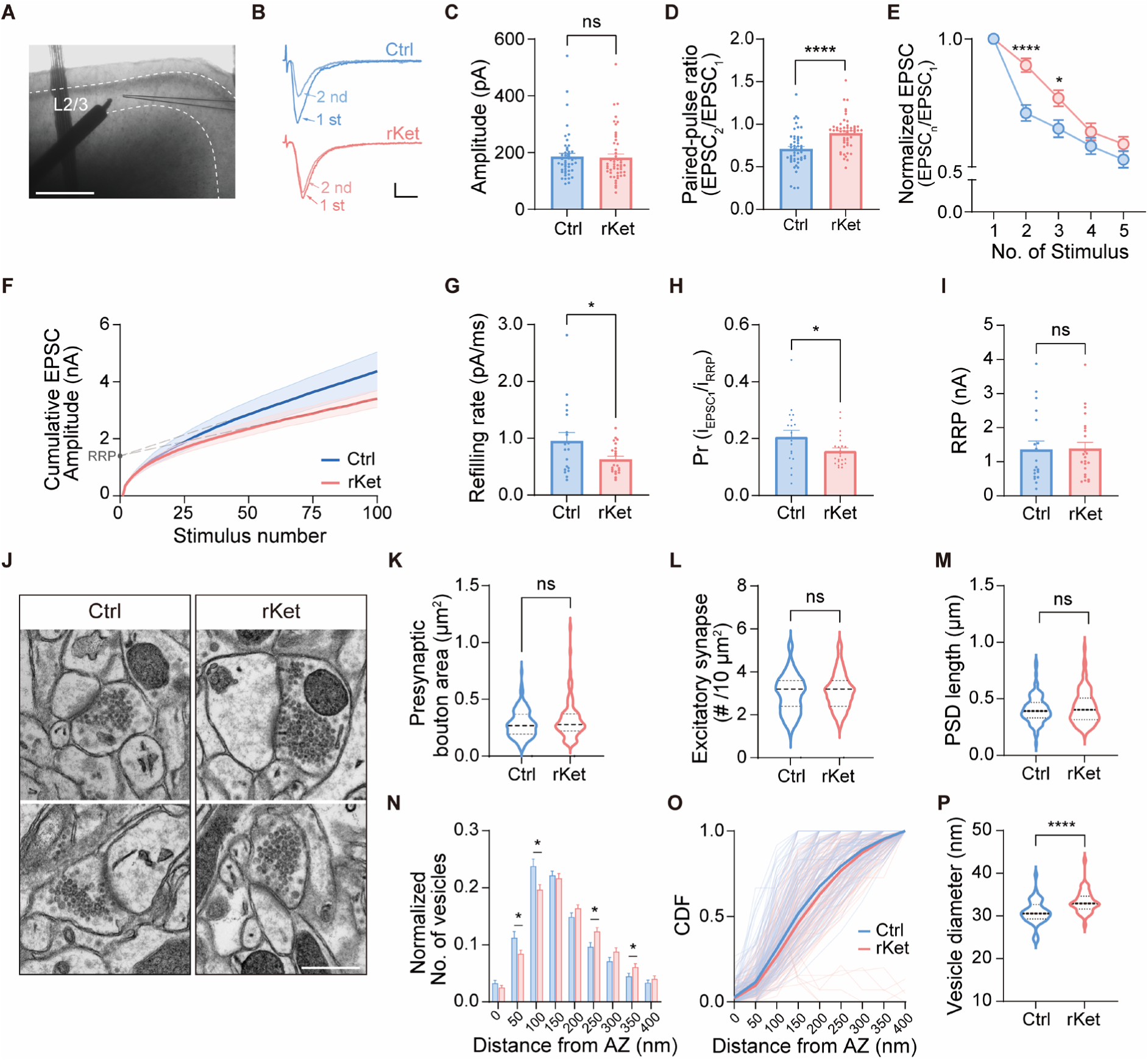
Release probability and vesicle organization of excitatory synapses in the dmPFC following chronic NMDAR hypofunction. **A–E,** Short-term plasticity analysis of evoked EPSCs using 10 Hz electrical stimulation (Ctrl: 49 cells from 12 mice; rKet: 51 cells from 12 mice). **A,** Representative coronal slice image showing the stimulating electrode positioned in layer 2/3 of the dmPFC during whole-cell recording. Recordings were performed in the presence of 100 µM picrotoxin. Scale bar: 500 μm. **B,** Representative EPSC traces. Scale bars: 10 ms, 50 pA. **C,** Average amplitude of the first evoked EPSCs, adjusted to approximately 180-200 pA. **D,** Paired-pulse ratio of evoked EPSCs at 500 ms interstimulus intervals. **E,** Evoked EPSCs during 10 Hz stimulation (1st to 5th), normalized to the first response. **F–I,** Analysis of synaptic release parameters using 33.33 Hz stimulus trains in the presence of 100 µM picrotoxin (Ctrl: 19 cells from 5 mice; rKet: 23 cells from 5 mice). **F,** Cumulative EPSC peak amplitude during 100 stimuli at 33.33 Hz. Summary graphs of **G,** refilling rate, **H,** readily releasable pool (RRP) size, and **I.** Probability of release (Pr). **J–P,** Ultrastructural analysis of excitatory synapses using transmission electron microscopy (TEM). **J,** Representative TEM images of the dmPFC layer 2/3 synapses. Scale bars: 500 nm. **K,** Presynaptic bouton area, **L,** Excitatory synapse density per 10 μm² of imaging area, **M,** Postsynaptic density (PSD) length, and **N,** Synaptic vesicle diameter at excitatory synapses. **O,** Cumulative distribution function (CDF) of vesicle distances from the AZ. Thin lines represent individual synapses; bold lines represent group averages. **P,** Diameter of synaptic vesicles. Data are presented as mean ± SEMs; \**p* < 0.05, \*\*\*\**p* < 0.0001, and ns, not significant by unpaired Student’s t-test compared with the Ctrl group.

To examine whether these functional deficits were accompanied by structural alterations, we conducted transmission electron microscopy of excitatory synapses in dmPFC layer 2/3. Gross synaptic morphology—including presynaptic bouton area, postsynaptic density (PSD) length, and synapse density—was unchanged between groups (Fig. 3K–M), suggesting that overall synaptic architecture was preserved. However, detailed vesicle analysis revealed nanoscale disruptions. Synaptic vesicles were positioned significantly farther from the active zone (AZ) in rKet mice, as indicated by a rightward shift in the cumulative distribution of vesicle distances (Fig. 3N–O). Additionally, vesicle diameter was significantly increased (Fig. 3P), suggesting altered vesicle dynamics or maturation. Together with confocal analysis (Extended Data Fig. 6), these findings indicate that the reduction in excitatory synaptic events cannot be explained by altered intrinsic excitability or changes in synapse number or morphology but instead arises from impaired Pr and vesicle refilling.

### Corticocortical and TF synaptic transmission in the dmPFC

To identify the input pathways responsible for reduced excitatory transmission onto dmPFC layer 2/3 pyramidal neurons, we investigated two major sources: bottom-up TF projections from the mediodorsal thalamus (MD) and local corticocortical (CC) excitatory connections within layer 2/3. Thalamocortical inputs provide direct feedforward excitation but are also amplified within the cortex by local recurrent excitatory circuits, whereby thalamorecipient pyramidal neurons recruit neighboring pyramidal cells^73,74^. Such recurrent excitatory connections have also been proposed as a cellular mechanism underlying persistent activity during STM^75–78^.

We first asked whether local CC connectivity was altered by chronic NMDAR hypofunction. To selectively activate layer 2/3 pyramidal neurons in the dmPFC, we used Wfs1-CreERT2 mice with tamoxifen-inducible expression of oChIEF, a stabilized, desensitization-resistant channelrhodopsin-2 (ChR2) variant. Optogenetic stimulation of layer 2/3 axons, paired with whole-cell recordings from neighboring pyramidal neurons, revealed no significant differences in optogenetically evoked EPSC (oEPSC) amplitude or short-term plasticity between rKet and control mice (Extended Data Fig. 7), indicating that CC connectivity is preserved.

We next assessed TF synapses, which are known to be critical for sustaining persistent activity and supporting STM^42,46–48^. oChIEF was selectively expressed in the MD to drive optogenetic stimulation of TF terminals while recording from dmPFC layer 2/3 pyramidal neurons. In contrast to the local CC inputs, rKet mice exhibited significantly reduced peak oEPSC amplitude and attenuated short-term depression (Fig. 4A–D), consistent with a reduction in Pr at TF synapses.

**Figure 4.**
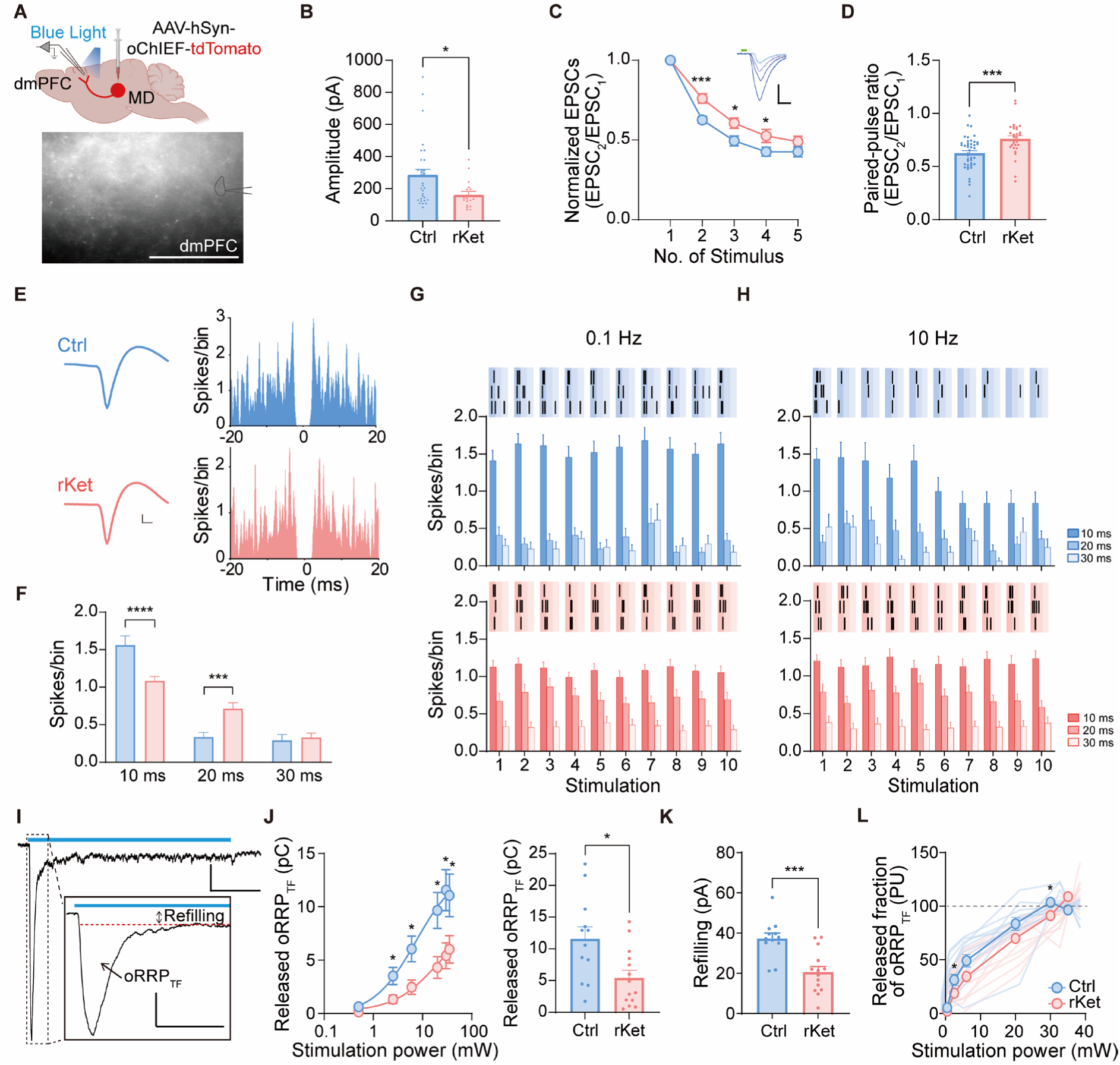
Chronic NMDAR hypofunction impairs TF synaptic transmission. **A–D,** Whole-cell recordings of optogenetically evoked thalamofrontal (TF) synaptic transmission (Ctrl: 32 cells, 11 mice; rKet: 17 cells, 7 mice). **A,** Schematic showing AAV-hSyn-oChIEF-tdTomato injection into the mediodorsal thalamus (MD) and whole-cell recordings from dmPFC pyramidal neurons. Representative image shows oChIEF-expressing MD axon terminals and the recorded neuron in dmPFC. Scale bar: 100 µm. **B,** Peak amplitude of the first optogenetically evoked EPSC (oEPSC) during 10 Hz stimulation. **C,** Normalized amplitudes of oEPSCs during a 10 Hz train. Inset, representative traces. Scale bars: 5 ms, 100 pA. **D,** Paired-pulse ratio of oEPSCs at a 500 ms interstimulus interval. **E–H,** Single-unit activity recorded in dmPFC during MD electrical stimulation. **E,** Representative waveforms (left) and autocorrelograms (right) of well-isolated single units in Ctrl and rKet mice. Scale bars: 500 µs, 10 µV. **F,** Average spike counts per bin within 10 ms, 20 ms, and 30 ms after each stimulation pulse (10 trials). **G, H,** Peri-stimulus histograms showing spike counts during **G.** 0.1 Hz and **H.** 10 Hz MD stimulation. **I–L,** Whole-cell recordings of optogenetically evoked readily releasable pool of TF synapses (oRRP_TF_) (Ctrl: 13 cells, 3 mice; rKet: 14 cells, 3 mice) in the presence of picrotoxin (100 µM), TTX (0.5 µM), and 4-AP (100 µM). **I,** Representative traces of synaptic currents evoked by 1 s optogenetic stimulation. The fast component represents vesicle depletion; the sustained component reflects vesicle refilling. Inset, expanded view of the fast component illustrating the oRRP_TF_ estimation; the red dashed line indicates the level used to quantify refilling. Scale bars: main trace, 250 ms, 50 pA; inset, 50 ms, 50 pA. **J,** Left, oRRP_TF_ charge plotted against stimulation power (log scale). Right, summary bar graph showing oRRP_TF_ at 30 mW (saturating intensity in controls). **K,** Summary bar graph of sustained refilling current amplitude at 30 mW, representing release of newly primed vesicles. **L,** oRRP_TF_ charge normalized to the maximum response per cell and expressed as pool units (PU), reflecting relative utilization of the vesicle pool. Maximum response was defined as the average oRRP_TF_ charge at 30–35 mW. Data are presented as mean ± SEMs. \**p* < 0.05, \*\**p* < 0.01, \*\*\**p* < 0.001, \*\*\*\**p* < 0.0001 and ns, not significant by unpaired Student’s t-test compared with Ctrl group.

To control for potential artifacts of ChR2 desensitization and to assess whether this synaptic impairment extends to the intact brain, we performed *in vivo* extracellular recordings in the dmPFC while electrically stimulating the MD. Firing responses were analyzed in 10 ms time bins from stimulus offset. At low-frequency stimulation (0.1 Hz), rKet mice showed a trend toward reduced firing in the initial 0–10 ms window, followed by a significant increase in the 10–20 ms bin (Fig. 4F). Although the overall spike count was similar between groups, the temporal distribution of responses was altered, indicating a loss of precise, time-locked activation. This shift suggests a degradation of feedforward inhibition, which normally sharpens excitatory response timing through rapid recruitment of local PV^+^ interneurons (Fig. 4G). During high-frequency stimulation (10 Hz), dmPFC neurons in control mice exhibited robust frequency-dependent suppression of spiking, whereas this suppression was blunted in rKet mice (Fig. 4H). Thus, the *in vivo* response pattern closely recapitulates the *ex vivo* findings of impaired TF synaptic dynamics under chronic NMDAR hypofunction.

Together, these *ex vivo* and *in vivo* findings identify TF synapses as a selective locus of presynaptic dysfunction under chronic NMDAR hypofunction, consistent with the STM deficits observed in rKet mice, although contributions from additional pathways cannot be excluded.

### Quantification of release parameters at TF synapses

To directly measure vesicle release capacity at TF synapses, we developed a modified optogenetic depletion assay. Traditional methods for estimating RRP size, such as high-frequency electrical or hyperosmotic challenge, lack spatial specificity and recruit mixed inputs. Optogenetic stimulation enables projection-specific activation of TF terminals, but strong optogenetic drive can evoke polysynaptic responses through recruitment of local recurrent circuits. To overcome this, we used prolonged (1 s) optogenetic stimulation of MD axon terminals while pharmacologically isolating monosynaptic responses with TTX and 4-AP, which block action potentials and enhance terminal depolarization, respectively (Fig. 4I–L). This protocol reliably evoked a biphasic current consisting of an initial rapid inward component followed by a sustained plateau. The fast component scaled with light intensity and saturated at ∼30 mW, followed by a prolonged component likely reflecting continuous vesicle recruitment. This pattern is reminiscent of the biphasic response evoked by hypertonic sucrose application, where the initial transient defines the RRP and the sustained component reflects ongoing vesicle refilling^79^.

We defined the optogenetically derived RRP at TF synapses (oRRP_TF_) as the transient charge released during the fast component of the biphasic response saturating stimulus intensity, estimated by subtracting the steady-state plateau from the total charge. oRRP_TF_ was significantly smaller in rKet mice, indicating a reduced pool of fusion-competent vesicles (Fig. 4J). Additionally, the amplitude of the sustained component — reflecting vesicle refilling — was significantly lower in rKet mice (Fig. 4K), consistent with the deficits in refilling rate observed in classical depletion experiments (Fig. 3G). To estimate release probability at TF synapses (Pr_TF_), we calculated the fractions of oRRP_TF_ released at each light intensity and compared these values between groups. The resulting input-output curve was right-shifted in rKet mice (Fig. 4L), indicating that equivalent stimulation released a smaller fraction of oRRP_TF_, consistent with a decrease in Pr (Fig. 3H).

Together, these results demonstrate that chronic NMDAR hypofunction selectively impairs TF synaptic transmission by reducing Pr, vesicle pool size, and refilling efficiency, raising the possibility that this projection represents a major site of synaptic vulnerability underlying STM impairment.

### Chemogenetic restoration of TF synaptic strength

Our findings so far collectively suggest that repeated ketamine exposure impairs STM performance and reduces synaptic efficiency in the dmPFC, particularly in the TF synapses. To test whether this synaptic dysfunction causally contributes to the observed behavioral deficits, we asked whether enhancing release efficacy at TF synapses could rescue STM performance. Prior studies indicate that Gq-coupled signaling pathways can facilitate presynaptic neurotransmitter release via activation of Munc13/PKC cascades^80^. Based on this, we employed a chemogenetic approach using hM3Dq, a Gq-coupled Designer Receptor Exclusively Activated by Designer Drugs (DREADD), to selectively potentiate TF synaptic transmission.

To selectively target MD → dmPFC projection neurons, we performed cholera toxin subunit B (CTB) conjugated with Alexa-488 retrograde labeling to identify MD neurons projecting to the dmPFC (Extended Data Fig. 8) and expressed hM3Dq in the corresponding MD region. This was achieved by injecting a Cre-dependent adeno-associated virus (AAV) encoding hM3Dq (AAV-DIO-hM3Dq) into the MD and a retrograde AAV-Cre into the dmPFC, ensuring selective expression in TF-projecting neurons. To activate hM3Dq, we administered compound 21 (C21), a selective agonist that engages Gq signaling. In parallel, we co-expressed oChIEF in MD neurons to allow optogenetic stimulation of TF axon terminals during whole-cell recordings from dmPFC layer 2/3 pyramidal neurons, enabling direct assessment of synaptic transmission following chemogenetic activation (Fig. 5A).

**Figure 5.**
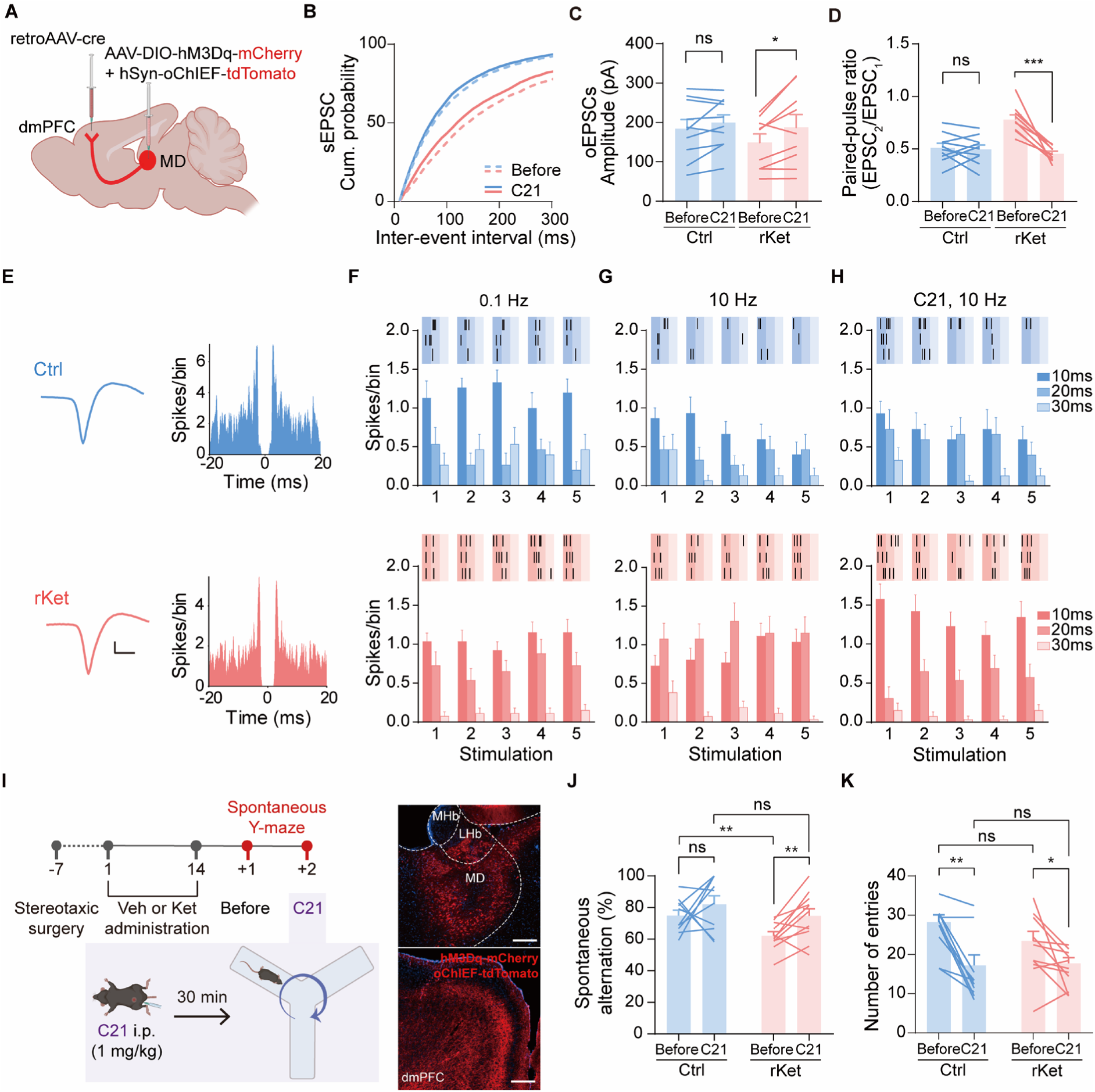
Chemogenetic activation of TF synapses restores synaptic function and STM performance in rKet mice. **A–D,** Whole-cell recordings from dmPFC pyramidal neurons examining the synaptic effects of Compound 21 (C21). **A,** Schematic of viral strategy. Cre-dependent hM3Dq-mCherry and Cre-independent oChIEF-tdTomato were co-injected into the MD, and retroAAV-Cre was delivered to the dmPFC, enabling chemogenetic activation and optogenetic monitoring of MD → dmPFC (TF) synapses. **B,** Cumulative probability distributions of spontaneous EPSC inter-event intervals recorded before and after bath application of C21 (1 μM) (Ctrl: 8 cells, 3 mice; rKet: 6 cells, 3 mice). Kolmogorov–Smirnov tests showed significant effects of C21 in both groups (\*\*\*\**p* < 0.0001). **C,** Peak amplitudes of oEPSCs during 0.1 Hz stimulation before and after C21 application. **D,** Paired-pulse ratios of oEPSCs at a 500 ms interstimulus interval (Ctrl: 10 cells, 3 mice; rKet: 9 cells, 3 mice). **E–H,** *i***n** *vivo* extracellular recordings of dmPFC single-unit activity during MD stimulation before and after systemic C21 administration (1 mg/kg, i.p.). **E,** Representative waveforms (left) and autocorrelograms (right) of well-isolated single units from Ctrl and rKet mice. Scale bars: 500 µs, 10 µV. **F, G,** Spike probability (spikes/bin) in response to MD stimulation at 0.1 Hz (F) and 10 Hz (G), averaged across the first five pulses. **H,** Spike responses to 10 Hz MD stimulation recorded ∼30 min after systemic C21 administration (binned at 10, 20, and 30 ms post-stimulus). **I–K,** Chemogenetic activation of MD → dmPFC projections during the spontaneous Y-maze task. **I,** Experimental timeline (left): AAV injections were performed in the MD and dmPFC, followed by 1 week of recovery and 14 days of daily vehicle or ketamine injections. Behavioral testing was conducted on two consecutive days: Day 1 (Before), baseline; Day 2 (C21), 30 min after C21 injection (1 mg/kg, i.p.). Representative fluorescence images (right) show hM3Dq-mCherry and oChIEF-tdTomato expression in the MD (scale bar: 200 µm) and dmPFC (scale bar: 100 µm), confirming targeted MD–dmPFC connectivity. LHb, lateral habenula; MHb, medial habenula. **J,** Spontaneous alternation performance before and after C21 administration. **K,** Total arm entries during the spontaneous alternation task. Data are presented as mean ± SEMs; within-group comparisons were analyzed using paired Student’s t-tests, and between-group comparisons using unpaired Student’s t-tests. \**p* < 0.05, \*\**p* < 0.01, \*\*\**p* < 0.001, and ns, not significant.

To validate the synaptic effects of chemogenetic activation, we performed *ex vivo* whole-cell recordings from layer 2/3 pyramidal neurons in the dmPFC. Bath application of C21 increased the frequency of sEPSCs in both control and rKet slices, consistent with enhanced presynaptic excitability and neurotransmitter release (Fig. 5B). To specifically assess TF synaptic efficacy, we measured optogenetically evoked EPSCs in response to oChIEF stimulation of MD terminals. In rKet slices, C21 application significantly increased oEPSC amplitude and restored the degree of short-term depression to levels comparable to those of control mice (Fig. 5C–D), indicating that Gq activation effectively enhanced presynaptic Pr_TF_. We next examined whether this synaptic enhancement was also evident *in vivo*. Single-unit activity was recorded in the dmPFC during electrical stimulation of the MD. Prior to C21 administration, control mice displayed robust frequency-dependent suppression of dmPFC firing during 10 Hz stimulation, whereas rKet mice showed attenuated suppression (Fig. 5F–G). Following C21 injection, dmPFC neurons in rKet mice exhibited restored firing suppression during 10 Hz MD stimulation (Fig. 5H), consistent with enhanced thalamic drive and recovery of TF synaptic dynamics observed in *ex vivo* recordings. Furthermore, the temporal profile of evoked responses was normalized, indicating a recovery of temporally precise thalamocortical integration (Fig. 5F–H).

### Chemogenetic enhancement of TF synaptic activity in STM task

Having established that chemogenetic activation restores TF synaptic transmission, we next asked whether this manipulation could rescue STM behavior. Spontaneous alternation performance in the Y-maze was assessed across two consecutive days: Day 1 without treatment, and Day 2 following systemic administration of C21 (30 minutes prior to testing; Fig. 5I). In rKet mice, spontaneous alternation significantly increased after C21 injection, reaching levels comparable to control animals (Fig. 5J), indicating restoration of STM function. While the total number of arm entries declined on Day 2 in both groups (Fig. 5K), likely due to habituation or reduced novelty-driven exploration^81^, this decrease did not account for the improvement in alternation rate, suggesting a genuine enhancement of memory performance.

Together, these results demonstrate that enhancing TF synaptic activity is sufficient to reverse STM impairments caused by NMDAR hypofunction, thereby establishing a causal link between TF synaptic dysfunction and cognitive deficits. This finding underscores the essential role of intact TF signaling in STM and identifies this pathway as a promising therapeutic target for treating NMDAR-related cognitive disorders.

### Discriminability of delayed activity during STM in the dmPFC

While our results so far demonstrated circuit-level TF dysfunction under chronic NMDAR hypofunction and its causal relationship with STM performance, it remained unclear how this impairment translates into prefrontal neural coding during STM performance. To test this, we examined task-related activity of dmPFC neurons during a Y-maze task, which requires mice to visit a previously unvisited arm—entailing two identical directional turns. We compared dmPFC activity at the delay zone between correct trials (Left–Left, LL and Right–Right, RR) and error trials (Left–Right, LR and Right–Left, RL) to determine whether the directional representation encoded by dmPFC population activity was diminished (Fig. 6A). We reasoned that in correct trials, mice successfully retained the memory of the first turn, whereas in error trials, the direction of the first turn was misremembered.

**Figure 6.**
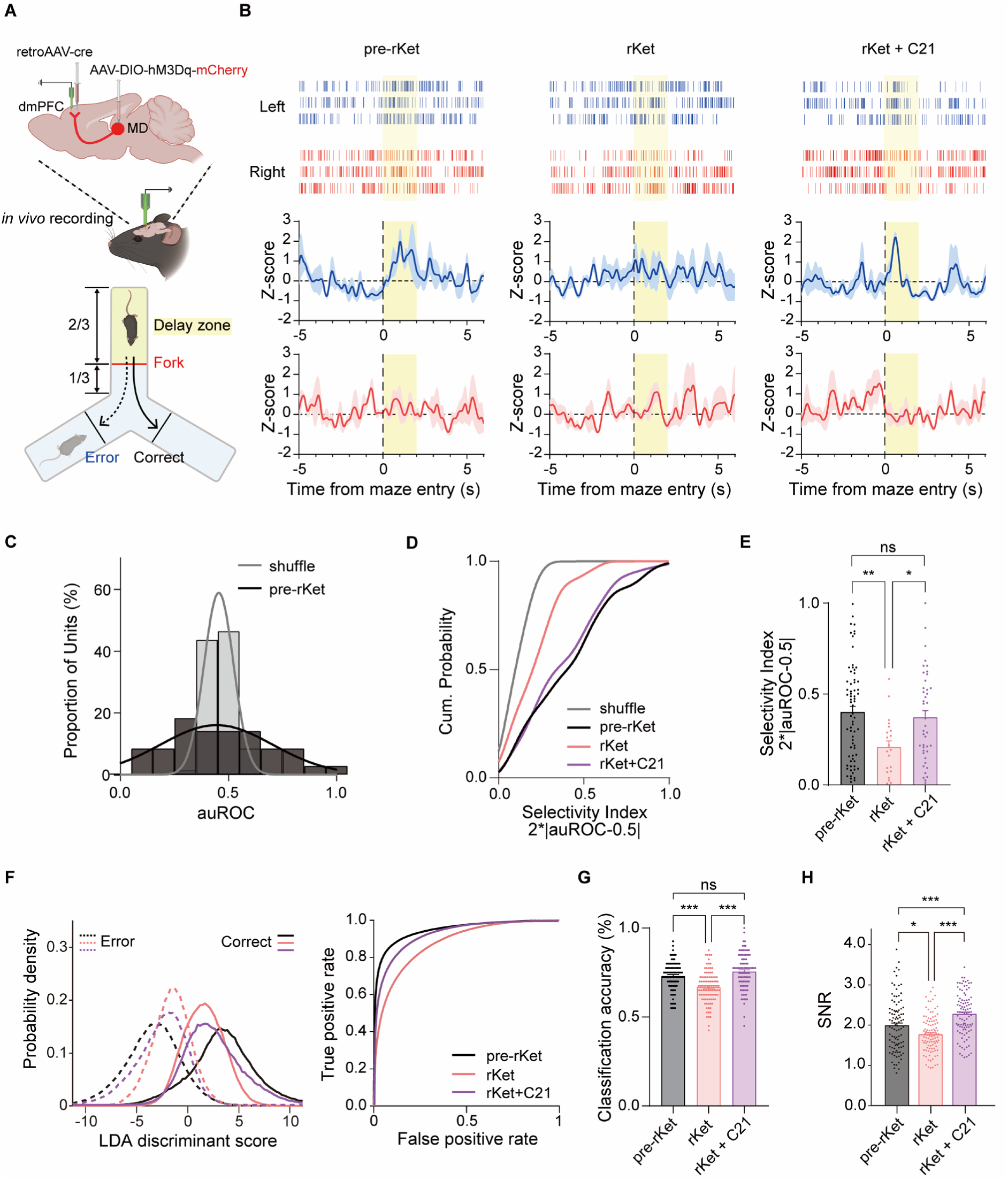
Reduced dmPFC firing and neural selectivity in repeated ketamine-treated mice are reversed by TF enhancement. **A,** Schematic of the experimental design for *in vivo* recording of thalamofrontal (TF) activity during the Y-maze task. **Top:** RetroAAV-Cre was injected into the dmPFC and AAV-DIO-hM3Dq-mCherry into the MD to express hM3Dq selectively in MD → dmPFC projection neurons. Recordings were made from dmPFC to monitor task-related activity. **Bottom:** Schematic of the Y-maze task assessing STM. Correct (solid) and error (dashed) trials were defined by whether the mouse repeated the same directional turn (LL or RR) or switched to the opposite arm (LR or RL) at the delay zone. **B,** Raster plots and peri-event time histograms of a representative dmPFC neuron during left (blue) and right (red) trials under pre-rKet, rKet, and rKet+C21 conditions, aligned to maze entry (time 0). The shaded yellow area indicates the delay zone (arm-entry to 2 s). **C–E,** Directional selectivity of dmPFC neurons based on area under the ROC curve (auROC). **C,** Distribution of auROC values within 2 s after arm entry for left vs. right trials (dark gray, pre-rKet; light gray, shuffled). **D,** Cumulative probability distributions of the selectivity index (2 × |auROC − 0.5|) for pre-rKet, rKet, and rKet+C21 conditions, with shuffled data shown for comparison. Kolmogorov–Smirnov tests indicated significant differences between pre-rKet and all other groups (*p* < 0.0001). **E,** Summary of mean selectivity indices (pre-rKet: 4 mice, 71 units; rKet: 4 mice, 22 units; rKet+C21: 4 mice, 43 units). Data are presented as mean ± SEM; one-way ANOVA with Bonferroni correction: \**p* < 0.05, \*\**p* < 0.01, ns, not significant. **F–H,** Population decoding analyses using linear discriminant analysis (LDA). **F,** LDA classifiers were trained on mean firing rates from 15 randomly selected single units in the delay zone to distinguish same-direction turns (LL, RR; correct) from opposite-direction turns (LR, RL; error). **Left**: Distributions of LDA scores for same-direction (solid) and opposite-direction (dashed) trials across pre-rKet (black), rKet (pink), and rKet+C21 (purple). **Right**: ROC curves derived from these scores. Kolmogorov–Smirnov tests revealed significant shifts across conditions (all *p* < 0.0001). **G,** Leave-one-neuron-out cross-validated LDA decoding of same-direction vs. opposite-direction turns. Forty iterations were averaged per accuracy estimate and bootstrapped 100 times to generate distributions. Bars show means; dots represent bootstrap iterations (\*\**p* < 0.001; ns, not significant). **H,** Population decoding of delay vs. fork activity using LDA. For each group, pseudo-populations were generated by randomly sampling neurons and trials across sessions. Discriminability was quantified as the signal-to-noise ratio (SNR) of LDA-projected scores (difference of means normalized by the pooled SD). Bars represent mean ± SEM; dots indicate bootstrap estimates. One-way ANOVA with Tukey’s post hoc test. One-way ANOVA with Tukey’s post hoc test: \**p* < 0.05, \*\**p* < 0.01, and ns, not significant.

We therefore recorded dmPFC neuronal activity across baseline (pre-rKet), rKet, and C21 conditions (Fig. 6B) to determine whether TF synaptic strength is reflected in cortical coding.

Specifically, we analyzed the direction selectivity of delay-period activity during the Y-maze task using receiver operating characteristic (ROC) analysis across the three conditions: baseline (pre-rKet), after rKet administration, and after C21 injection (rKet + C21). Direction selectivity was quantified using a selectivity index, calculated as twice the difference between the area under the receiver operating curve (auROC) value and that of a random discriminator (see Methods). In correct STM task trials of the baseline condition, the selectivity indices of recorded neurons were significantly higher than those obtained from shuffled data. Consistent with the idea that reduced STM performance under rKet is associated with diminished neural coding in dmPFC neurons, discriminability was significantly decreased by rKet and restored following enhancement of TF synapses with C21 (Fig. 6C–E).

We then analyzed whether STM-dependent population coding in the dmPFC was disrupted by repetitive NMDAR hypofunction using linear discriminant analysis (LDA). To control for differences in the number of recorded units across trials, we pooled all recorded units from all sessions and employed a subsampling-based classification approach. Specifically, we randomly selected 15 units to train a LDA classifier that best distinguished correct from error population activity. We then assessed classification performance by calculating the area under the receiver operating characteristic (ROC) curve using a separate set of 15 randomly selected units. This procedure was repeated 1,000 times to obtain a distribution of decoding performance across balanced subsamples.

Compared to the ROC values obtained during pre-injection, rKet administration led to a significant reduction in ROC, indicating that dmPFC activity during correct trials became more similar to that of incorrect trials (Fig. 6F, pink). This suggests a degradation of direction-selective population coding under NMDAR hypofunction. Notably, chemogenetic strengthening of TF synaptic transmission restored the distinction between correct and incorrect trials, as reflected by a significant recovery of ROC values (Fig. 6F, purple), consistent with the idea that the disrupted population coding is due to the weakening of TF synapses.

Because of the limited number of recorded units and trials, the subsampling approach could potentially over-represent specific subsets of neurons. To mitigate this issue, we employed two complementary strategies. First, without subsampling, we implemented a leave-one-neuron-out cross-validation procedure: an LDA classifier was iteratively trained using all but one unit to distinguish correct from incorrect trial activity and tested on the held-out unit. Consistent with the results in Fig. 6F, this analysis revealed reduced prediction accuracy in the rKet group compared to the pre-Ket condition, which was restored following C21 administration (Fig. 6G). Second, we compared dmPFC population activity between the delay zone and the fork region. We reasoned that in the delay zone, mice must retain directional information from the previous turn, whereas at the fork, they initiate the subsequent action based on that memory. Thus, if directional information is successfully maintained, population activity in the delay zone should be distinguishable from that at the fork. This analysis also provided greater statistical power, as it allowed inclusion of a larger number of trials independent of behavioral outcome. Similarly, rKet administration significantly reduced the signal-to-noise ratio (SNR) of dmPFC population activity, an effect that was reversed by chemogenetic enhancement of TF synaptic strength (Fig. 6H).

Together, these results demonstrate that TF synaptic efficacy governs the fidelity of STM-dependent coding in dmPFC during STM, linking circuit-level disruption under NMDAR hypofunction to impaired population-level representations.

## Discussion

Weakening of the TF connection has been consistently observed in patients with schizophrenia, and several studies have reported that the strength of this connection correlates with symptom severity^53,58^. However, these findings have remained correlative.

In this study, we identified a projection-specific presynaptic mechanism underlying cognitive deficits induced by chronic NMDAR hypofunction. Mirroring the clinical observations in schizophrenia, repeated ketamine exposure selectively weakened TF synaptic transmission onto layer 2/3 pyramidal neurons in the dmPFC, characterized by reduced Pr, a smaller RRP, and diminished vesicle refilling efficiency. This synaptic weakening was accompanied by impaired population-level coding in the dmPFC during STM performance, as evidenced by reduced direction-selective decoding accuracy. Furthermore, chemogenetic enhancement of TF synaptic transmission was sufficient to restore both synaptic efficacy and STM-related neural coding, as well as behavioral performance in rKet-treated mice. These findings suggest that TF circuit dysfunction may represent both a mechanistic substrate for STM deficits and a potential therapeutic target in schizophrenia.

### Homeostatic regulation of presynaptic release efficiency: contrast with E–I imbalance

These findings extend current models of NMDAR hypofunction in schizophrenia, which have largely emphasized E-I imbalance due to preferential impairment of fast-spiking interneurons^68^. While acute NMDAR antagonism produces cortical disinhibition and hyperexcitability^68,82–84^, the hypofunction observed in schizophrenia reflects a chronic state^7–9^. Our data demonstrated that chronic NMDAR hypofunction, which more closely mimics the pathophysiology of schizophrenia, instead weakens excitatory drive at TF synapses (Fig. 4C). We interpret this as a form of homeostatic synaptic plasticity, whereby cortical circuits adjust parameters regulating synaptic strength and excitability to stabilize activity under sustained disruption of excitatory drive^85,86^. Typically, homeostatic regulation is thought to occur postsynaptically, through changes in receptor strength or intrinsic excitability^87,88^. In our study, we observed an increased AMPA–NMDA ratio, consistent with prior reports^86^ that NMDAR blockade can upregulate AMPAR surface expression (Extended Data Fig. 9). However, despite such postsynaptic scaling, we found reduced synaptic gain, as shown by our input–output analysis of fEPSPs (Extended Data Fig. 10). Furthermore, intrinsic excitability was unchanged (Fig. 2), ruling out compensatory changes in postsynaptic excitability. Instead, we found convergent evidence for reduced presynaptic efficiency in rKet mice, including attenuated short-term depression, decreased Pr, and diminished vesicle availability and refilling (Fig. 3), consistent with previous reports of impaired presynaptic neurotransmitter release^89–91^.

It is notable that presynaptic downregulation is not a pan-cortical phenomenon. Release efficiency of somatosensory thalamocortical synapses was not affected by repeated ketamine administration (Extended Data Fig. 5). We believe vulnerability of the TF synapses may be attributed to the delayed maturation of the prefrontal circuits^21^. Repeated ketamine administration leads to sustained NMDAR blockade, likely leading to prolonged excitability elevation of pyramidal neurons. Notably, the juvenile to adolescent period, which coincides with the timing of our drug administration, represents a robust phase of synaptic pruning and refinement in the dmPFC^24^, driven in part by complement-mediated synapse pruning where microglia engage in phagocytic engulfment and elimination^92,93^. This developmental window is also marked by refinement of proper synaptic contacts between the MD and PFC and axonal pruning in local PFC circuits^22^. Critically, this window overlaps with the typical age of onset for schizophrenia, suggesting that perturbations to TF circuit maturation during adolescence—such as those induced by chronic NMDAR hypofunction—could contribute to disease vulnerability. Our findings thus provide a potential mechanistic link between altered developmental synaptic refinement and the emergence of prefrontal dysfunction in early-stage psychosis.

### Speculative molecular mechanisms of presynaptic downregulation

Although elucidating the molecular basis of these presynaptic changes is beyond the scope of the present work, several postmortem studies in schizophrenia provide relevant clues. Abnormal protein–protein interactions within the presynaptic release machinery have been reported, including disrupted binding among SNARE proteins and increased affinity of syntaxin-1 for Munc18-1 and complexin in orbitofrontal and anterior cingulate cortices^94^. Munc18-1 is classically thought to stabilize the “closed” conformation of syntaxin-1, preventing productive SNARE assembly^95,96^, whereas the “open” conformation lowers the energy barrier for membrane fusion and accelerates replenishment of the readily releasable pool^97^. Complexin also plays a modulatory role in fusion^98,99^. Thus, excessive binding affinity to these regulatory proteins could reduce the efficiency of vesicle release and replenishment. In line with this, phosphorylated syntaxin-1 is reduced in schizophrenia brain tissue, which has been suggested to impair syntaxin-1 binding to its protein partners^94^. Within this framework, the enhanced Munc18–SNARE association described in schizophrenia provides a plausible molecular explanation for the reduced Pr and refilling we observed at TF synapses under chronic NMDAR hypofunction.

Our findings appear somewhat contradictory to a recent study reporting that chronic phencyclidine administration leads to reduced PV^+^ interneurons activity and consequently increased activity in the prelimbic cortex, which was proposed to causally impair working memory^28,29^. The variability reported across studies employing pharmacological NMDAR hypofunction^28,29^ may stem from age-dependent differences in the underlying homeostatic plasticity mechanisms^100,101^.

### Potential mechanism of STM deficits

Although the MD is critical for sustaining PFC activity during STM^42,46,102^ and its disruption is linked to psychiatric disorders with cognitive deficits^60^, the precise mechanisms underlying this contribution remain unclear. One possibility is that MD–PFC synapses convey mnemonic content through recurrent TF loops. Supporting this idea, recordings in nonhuman primates have shown that many MD neurons exhibit task-relevant directional selectivity during working memory tasks^103^, suggesting that MD inputs can transmit specific mnemonic information to the PFC. Our findings provide circuit-level support for this perspective. We observed that reduced TF synaptic strength under chronic NMDAR hypofunction was associated with diminished discriminability of dmPFC neurons during STM, whereas chemogenetic strengthening of TF inputs restored discriminability along with behavioral performance (Fig. 5-6). These results demonstrate that TF efficacy directly constrains cortical coding capacity, establishing a causal link between TF synaptic dysfunction, impaired neural representations, and STM deficits.

Alternatively, the MD–PFC pathway may regulate the flexible allocation of attention and cognitive resources, rather than encoding mnemonic content directly. In this framework, MD activity is indispensable for sustaining PFC selectivity and STM performance by enhancing functional connectivity among PFC neurons without necessarily increasing their firing rates^48^. Notably, Schmitt and colleagues found that enhanced MD activity selectively increases PV^+^ interneuron firing, consistent with our observation of reduced TF feedforward inhibition (Fig. 4F). It is conceivable that reduced feedforward inhibition arises from both weakened TF input strength and a decrease in the number of PV^+^ interneurons, leading to a diminished capacity for timely inhibitory control. The temporal window of excitatory integration is regulated by PV^+^ interneurons in both sensory cortices^104^ and higher-order areas such as the PFC^105,106^. While the importance of temporal precision in higher-order cortical processing is less established than in sensory systems, our recent findings in the posterior parietal cortex indicate that temporal inaccuracy in sequential delay activity predicts task errors^107^. This raises the possibility that weakened TF drive diminishes feedforward inhibition, broadens the temporal integration window, and thereby disrupts the precision of activity patterns essential for STM. Consistent with the idea that TF input contributes to stabilizing task-relevant representations, we found that STM-related population coding in the dmPFC was weakened under rKet and partially restored by TF synaptic strengthening (Figs. 5G vs. 5H). A more systematic test of this hypothesis will require directly manipulating feedforward inhibition independently of thalamic input strength, while assessing the reliability of sequential activity during STM.

### Limitations of the study

Several limitations warrant further investigation. First, although chemogenetic activation of MD → PFC projections restored STM performance, the durability of this effect remains unknown. Given prior evidence that thalamic input can drive cortical plasticity^108^, it will be important to determine whether repeated stimulation produces lasting structural or functional adaptations. Second, our study focused on chronic NMDAR hypofunction during adolescence, a critical window for PFC maturation. It remains to be determined whether similar circuit dysfunctions arise when NMDAR signaling is disrupted in adulthood, such as under conditions of substance abuse or late-onset psychiatric illness. Defining the temporal window of NMDAR-dependent vulnerability will be essential for understanding how developmental versus acquired disruptions of this pathway contribute to cognitive deficits.

## Conclusion

In conclusion, our findings demonstrate that chronic NMDAR hypofunction during adolescence impairs STM and reduces synaptic efficiency in the dmPFC, primarily through diminished Pr_TF_. By showing that chemogenetic enhancement of TF connectivity rescues both synaptic efficacy and behavioral performance, we identify this pathway as a critical and modifiable circuit locus of cognitive vulnerability. Targeting TF connectivity may therefore represent a promising therapeutic strategy for schizophrenia, and other neuropsychiatric conditions characterized by prefrontal dysfunction.

## Methods

### Animals

All experimental procedures involving the use of live mice or their tissues were conducted in accordance with the Korea Brain Research Institute guidelines and were approved by the Animal Care and Use Committee (KBRI IACUC No. IACUC-23-00046 and IACUC-24-00019). Drug administration was initiated in three-to four-week-old male mice. C57BL/6NHsd mice, originating from the National Institutes of Health, were purchased from KOATECH (South Korea). For PV^+^ cell counting, PV-tdTomato mice were generated by crossing homozygous B6.129P2-B6.129P2-Pvalb^tm1^(cre)^Arbr^/J (stock #017320, The Jackson Laboratory, USA) with homozygous B6.Cg-Gt(ROSA)26Sor^tm9(CAG-tdTomato)Hze^/J (stock #007909, The Jackson Laboratory, USA). Wfs1-Tg3-CreERT2 mice (stock #009103, The Jackson Laboratory, USA) were used to enable layer 2/3-specific optogenetic stimulation in the dmPFC. To induce Cre recombination, mice were fed a tamoxifen-containing diet (500 mg/kg tamoxifen USP; TD.130857, Envigo, USA). Animals were given mixed chow for 2 days to allow acclimation, followed by exclusive access to the tamoxifen diet for 5 consecutive days after AAV injection, with daily intake not exceeding 80 mg tamoxifen per kg body weight. Mice were maintained on a 12-h light/dark cycle with food and water available *ad libitum*, and the breeding room was kept at 22 ± 2 °C with a relative humidity of 50 ± 10%.

### Systemic NMDAR antagonist administration

Ketamine hydrochloride (Yuhan Corporation, South Korea) was dissolved in sterile saline (0.9% NaCl) for injection. Mice received intraperitoneal (i.p.) injections of vehicle or ketamine (30 mg/kg) at a volume of 10 ml/kg once daily for 5 or 14 consecutive days.

### Electrophysiology

#### Slice preparation

Acute slices were prepared from five-to six-week-old male C57BL/6NHsd mice. Transcardial perfusion was performed under deep anesthesia induced by sodium pentobarbital (70 mg/kg, i.p.). Brains were rapidly removed and immersed in ice-cold cutting solution containing the following (in mM): 110 choline chloride, 2.5 KCl, 25 NaHCO_3_, 1.25 NaH_2_PO_4_, 25 Glucose, 0.5 CaCl_2_, 7 MgCl_2_·6H_2_O, 11.6 Sodium L-ascorbate, and 3 Sodium pyruvate. Thereafter, 300 μm-thick coronal slices were prepared using a vibratome (VT1200S, Leica Biosystems, Germany). The slices were then incubated for 30 min at 32 °C in artificial cerebrospinal fluid (aCSF) containing the following (in mM): 119 NaCl, 2.5 KCl, 26 NaHCO_3_, 1.25 NaH_2_PO_4_, 20 Glucose, 2 CaCl_2_, 1 MgSO_4_, 0.4 L-Ascorbic acid, and 2 Sodium pyruvate. During sectioning, the solutions were saturated with carbogen (95% O_2_, 5% CO_2_). Osmolality was adjusted to 306 ± 2 mOsm. For measuring IPSCs, aCSF contained 126 NaCl, 3.5 KCl, 25 NaHCO_3_, 1 NaH_2_PO_4_, 20 Glucose, 1 CaCl_2_, 0.5 MgSO_4_, 0.4 L-Ascorbic acid, and 2 Sodium pyruvate to a final pH of 7.4. Osmolality was adjusted to 312 ± 2 mOsm. All solutions were saturated with carbogen (95% O_2_, 5% CO_2_) to a final pH of 7.4.

#### Whole-cell recordings

The slices were transferred to a submerged recording chamber continuously perfused with aCSF saturated with carbogen (95% O₂, 5% CO₂) for whole-cell recordings. Layer 2/3 neurons in the dmPFC were visualized using an upright microscope (BX51WI, Olympus, Japan) equipped with differential interference contrast optics and a 40X water-immersion objective (NA 0.8, Olympus, Japan). All recordings were performed at 30 ± 2 °C, with fresh aCSF perfused at a rate of ∼3.6 mL/min. Glass electrodes were pulled from borosilicate glass capillaries (G150TF-4, Warner Instruments, USA) using a pipette puller (P-1000, Sutter Instruments, USA) to yield a tip resistance of 3.5–5 MΩ. For EPSCs recordings, the internal solution contained (in mM): 120 potassium gluconate, 10 KCl, 10 HEPES, 4 NaCl, 10 Na₂-phosphocreatine, 4 MgATP, 0.3 NaGTP, and 0.2 EGTA (pH 7.2; 296 ± 2 mOsm). For spontaneous IPSCs recordings, the internal solution contained (in mM): 120 KCl, 10 HEPES, 4 MgATP, 0.3 NaGTP, 10 Na₂-phosphocreatine, and 3 QX-314 bromide (pH 7.2; 290 ± 2 mOsm). For NMDA/AMPA ratio measurements, the internal solution contained (in mM): 125 CsMs, 5 CsCl, 10 HEPES, 0.2 EGTA, 4 MgATP, 0.3 NaGTP, 10 Na_2_-phosphocreatine, and 3 QX-314 bromide (pH 7.2; 295 ± 2 mOsm). Electrophysiological signals were recorded using a MultiClamp 700B amplifier and digitized with a Digidata 1550A interface (Molecular Devices, USA) at a sampling rate of 10 kHz. For EPSC measurements, signals were low-pass filtered at 1 kHz. Data were acquired using Clampex 10.5 software (Molecular Devices, USA).

Spontaneous synaptic transmission was measured in neurons voltage-clamped at-70 mV, and miniature excitatory postsynaptic currents were recorded under the same holding potential in the presence of 0.5 µM TTX. To evoke synaptic responses in the dmPFC, a concentric bipolar stimulation electrode (30214, FHC Inc., USA) was positioned in layer 2/3 near the recorded cell. Electrical stimulation was delivered at 0.1 Hz to assess basal transmission, at 10 Hz to evaluate short-term plasticity, and at 33.33 Hz in the dmPFC or 50 Hz in the S1 to deplete the RRP of vesicles. Stimulus pulses had a duration of 300 µs and were kept below 5 V, and recordings were taken at sites exhibiting stable, predominantly monosynaptic EPSCs.

For NMDA/AMPA ratio measurements, 100 µM picrotoxin was bath-applied throughout the recording to block GABA_A_ receptor-mediated currents. AMPAR-mediated currents were defined as the peak amplitude at a holding potential of –70 mV, whereas NMDAR-mediated currents were quantified as the mean amplitude over a 20 ms window beginning 47 ms after stimulus onset at 0 mV, a time window chosen to ensure minimal contamination from AMPAR-mediated responses. Paired-pulse ratio was calculated from 10 successive stimuli delivered at 10 Hz, as the ratio of the second EPSC amplitude to the first.

Optogenetic stimulation was delivered near the recorded cell at sites that produced stable, predominantly monosynaptic currents. Light (470 nm) was delivered through a 200 µm core diameter optic fiber (0.22 N.A.; Doric Lenses, Canada) positioned above the slice. The light pulse duration was adjusted between 2 and 10 ms to achieve reliable responses, and for prolonged (1 s) stimulation of MD axon terminals, light intensity was increased up to 35 mW. To isolate monosynaptic responses, 0.5 µM TTX, 100 µM 4-AP, and 100 µM picrotoxin were bath-applied. The fast component of the evoked response was defined as the optically induced readily releasable pool of TF synapses (oRRP_TF_), and the amplitude of the sustained component was defined as the vesicle refilling response.

#### Field EPSP recordings

Field EPSPs (fEPSPs) were recorded using 2–3 MΩ glass pipettes filled with standard aCSF and positioned in layer 2/3 of the dmPFC. Electrical stimuli (100 µs duration) were delivered at 0.05 Hz through a bipolar concentric electrode (TM33CCINS, WPI, USA) placed in layer 5. Stimulus intensity was determined by constructing an input–output curve in which the fEPSP slope was plotted against stimulus strength. For each intensity, three consecutive responses were evoked at 0.05 Hz and averaged for analysis.

#### Quantification of synaptic responses

Spontaneous and miniature EPSCs/IPSCs were detected using a template-matching algorithm in Clampfit from 1-min recording segments and subsequently filtered with custom MATLAB scripts. Events with amplitudes ≤ 8 pA, time-to-peak ≥ 10 ms, or inter-event intervals ≤ 5 ms were excluded from analysis. Identical detection and filtering criteria were applied across all groups to minimize electrical noise and exclude events that were read as overlapping.

For estimation of the RRP and vesicle refilling rate, trains of 100 stimuli were delivered at 33.33 Hz for dmPFC recordings and 50 Hz for S1BF recordings to deplete synaptic responses. The cumulative amplitude of evoked EPSCs during the stimulation train was plotted, and the linear portion of the late steady-state phase was extrapolated to the y-intercept to estimate the RRP (pA). The slope of this linear fit was defined as the vesicle refilling rate (pA/ms).

### Stereotaxic Viral Injection

All stereotaxic injections were performed using a stereotaxic frame (RWD Life Science, USA), and body temperature was maintained with a heating pad (FHC, USA). To prevent corneal drying, a thin layer of Vaseline (Unilever, USA) was applied to the eyes. Viral or tracer injections were delivered through a Hamilton microsyringe (32–35 gauge tip) connected to an UltraMicroPump (WPI, USA) and controlled by a Micro4 controller (WPI, USA) at a rate of ∼50 nL/min.

Mice were bilaterally injected with retroAAV-EF1a-Cre (2.5 × 10¹³ genome copies per mL (GC/mL), Addgene #55636-AAVrg) or retroAAV-EF1a-mCherry-IRES-Cre (7.0 × 10¹² GC/mL, Addgene #55632-AAVrg) into the dmPFC layer 2/3 (300 nL per site), and with AAV2-hSyn-DIO-hM3Dq-mCherry (2.0 × 10¹³ /mL, packaged by VIUS from Addgene plasmid #44361) into the MD (300 nL per side). AAV1/2-hSyn-oChIEF-mCherry-WPRE (1.0 × 10¹³ GC/mL, Sirion Biotech GmbH #KA16140_AA3929) was also injected bilaterally into the MD (200 nL per side), either alone or in combination with hM3Dq for electrophysiological experiments. For layer-specific targeting, AAV2/5-EF1a-DIO-oChIEF(E163A/T199C)-P2A-dTomato-WPRE-bGH polyA (2.6 × 10¹² GC/mL, BrainVTA # PT-0228) was bilaterally injected into the dmPFC (200 nL per side) of Wfs1-CreER mice. For projection labeling, AAV5-EF1a-Cre (4.4 × 10¹² GC/mL, UNC Vector Core) and AAV5-hSyn-FLEX-mGFP-2A-Synaptophysin-mRuby (4.1 × 10^13^ GC/mL, VectorBuilder #P210527-1012euv) were mixed at a 1:1 ratio and bilaterally injected into the MD (200 nL per side).

Coordinates were as follows: dmPFC, AP 1.50–1.60 mm from bregma, ML ±0.25 mm from midline, DV 1.30–1.10 mm from the skull surface; MD, AP 1.00 mm from bregma, ML ±0.40 mm from midline, DV 3.50–3.10 mm from the skull surface. Coordinates were based on the atlas of Paxinos and Franklin^109^. After completion of the injection, the needle was left in place for at least 5 min to allow viral diffusion, after which it was slowly retracted. The scalp incision was then sealed using a tissue adhesive, Vetbond (084-1469SB, 3M, USA). Animals were allowed to recover on a heating blanket before being returned to their home cages and received postoperative analgesia with ketoprofen (5 mg/kg, subcutaneous, once daily for two days).

### Histological analysis

For PV^+^ cell quantification, we used PV-tdTomato mice generated by crossing homozygous B6.129P2-Pvalb^tm1(cre)Arbr^/J (stock #017320, The Jackson Laboratory) with homozygous B6.Cg-Gt(ROSA)26Sor^tm9(CAG-tdTomato)Hze^/J (stock #007909, The Jackson Laboratory). For histological experiments, four control (2 males, 2 females) and four repetitive ketamine–treated (rKet; 2 males, 2 females) mice were used. During postnatal week 3, mice received drug administration for two consecutive weeks. On the day following the final administration, mice were anaesthetized with sodium pentobarbital (70 mg/kg, i.p.) and transcardially perfused with 10 mL phosphate-buffered saline (PBS; BPB-9104, Tech & Innovation, South Korea) followed by 10 mL ice-cold 4% paraformaldehyde. Dissected brains were post-fixed in 4% paraformaldehyde for 24 h at 4 °C and then transferred to 30% sucrose in PBS for cryoprotection. After 2 days at 4 °C, brains were embedded in frozen section compound (3801480, Leica Biosystems, Germany) and coronally sectioned at 40 μm using a cryostat (CM1860, Leica Biosystems, Germany). Sections were washed in PBS and mounted on slides with DAPI-containing mounting medium (H-1200, Vector Laboratories, USA).

Coronal brain sections were cut at 40 μm thickness, and every fourth section (160 μm interval) was collected, yielding six sections spanning from +1.94 mm to +1.18 mm relative to bregma. Slides were scanned using a digital slide scanner (Pannoramic SCAN II FL, 3DHISTECH, Hungary). Cell counting was performed in ImageJ/Fiji by manually identifying DAPI-positive nuclei and tdTomato-positive PV^+^ interneurons. Quantification was limited to the medial prefrontal cortex, specifically the prelimbic (PrL), infralimbic (IL), and cingulate cortex areas 1 (Cg1) and 3 (Cg3).

To identify the precise location of thalamic neurons in the MD that project to the dmPFC—where our *ex vivo* recordings were conducted—we utilized CTB–Alexa 488 (Extended Data Fig. 8).

Confocal images were acquired as Z-stacks with an optical step size of 0.5 μm using 10× and 60× objectives on a super-sensitivity NIR confocal microscope (STELLARIS 8, Leica Microsystems, Germany). Image acquisition parameters (laser power, detector gain, and offset) were kept constant across all slices. Image analysis was performed in ImageJ/Fiji by generating sum-intensity projections from 30 optical sections. Regions of interest (ROIs) were manually defined, and fluorescence intensity was quantified as the raw integrated density divided by ROI area (a.u./μm²).

### Chemogenetic manipulations

To manipulate MD → dmPFC projection neurons, AAVs were injected into layer 2/3 of the dmPFC and the MD of three-week-old mice. Behavioral testing was conducted six weeks after viral injection, followed by slice recordings from the same animals upon completion of behavioral experiments. For behavioral procedures, mice were allowed to recover for 3 weeks, during which they received daily drug administration during the final 2 weeks. For *in vivo* hM3Dq-mediated activation, mice received i.p. injections of the DREADD agonist C21 (1 mg/kg; HB4888, Hello Bio, UK) at a volume of 10 mL/kg. C21 was used instead of clozapine-N-oxide (CNO) because it does not metabolize into clozapine or CNO and exhibits higher receptor specificity and longer-lasting central activity^110,111^. In the behavioral rescue experiments (Fig. 5), C21 was administered 30 min before testing to transiently activate MD → dmPFC projection neurons.

### Behavioral tests

#### Open field test

Locomotor activity of mice was evaluated using the open field test in polystyrene enclosures (40 × 40 × 40 cm). Mice were placed in the center of the box and videotaped individually. The center area was defined as the central 20 × 20 cm zone. Tracking was analyzed using SMART 3.0 Video Tracking Software. Total distance traveled and time spent in the center area were analyzed.

#### Custom-designed Y-maze task

A custom-designed Y-maze equipped with motorized doors was used to precisely control arm access and monitor gating times during each task phase. During the 14-day administration period, mice underwent food restriction on days 9–11, with body weight maintained at 70–80% of their weight on day 8. On day 11, mice were allowed to freely explore the Y-maze for ∼20 min to familiarize themselves with the environment. Training began on day 12. In each trial, during the sample phase, one of the three arms was opened while the other two were blocked by motorized doors, forcing the mouse to enter the accessible arm to obtain a sucrose pellet placed at the end. Once the mouse fully entered the arm, its door closed, and one of the previously blocked arms opened, initiating a 30-s delay phase during which the mouse was confined. After the delay, the sample arm door was reopened, allowing the mouse to choose between the previously visited and the newly opened arm. A correct choice—entering the previously unvisited arm—was rewarded with another sucrose pellet. Each day comprised two sessions (morning and afternoon), with 10 trials per session, conducted over two consecutive days (days 12–13). The sequence of target arms was randomized to prevent consecutive trials requiring the same directional choice. Performance was assessed on day 14.

#### Spontaneous Y-maze test

To further assess STM, we conducted the spontaneous alternation task, which does not involve reward-based learning. Mice were placed in a standard Y-maze and allowed to explore freely for 8 minutes. All arm entries were recorded, and spontaneous alternation was calculated as the percentage of alternations using the following formula:

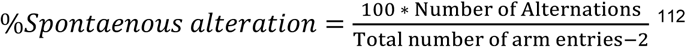

Tracking was analyzed using SMART 3.0 video-tracking software. Time spent in the middle zone (%) was analyzed.

### Electron microscopy (EM)

A separate group of mice, age-matched to those used for electrophysiological experiments (Ctrl: 4 mice; rKet: 4 mice) were deeply anesthetized and perfused transcardially with 0.15 M cacodylate buffer (pH 7.4) containing 2% paraformaldehyde and 2.5% glutaraldehyde. Brains were removed, and 150 μm coronal sections were cut on a vibratome (VT1200S, Leica Biosystems, Germany). Sections encompassing layer 2/3 of the dmPFC were dissected into small tissue blocks for EM processing.

For synaptic vesicle analysis, blocks were rinsed in 0.15M cacodylate buffer and post-fixed for 1 hour in 2% osmium tetroxide/1.5% potassium ferrocyanide. After distilled water washes, samples were incubated in 1% thiocarbohydrazide (223220, Sigma-Aldrich, USA) for 20 minutes, followed by a second 2% osmium tetroxide treatment for 30 minutes. Specimens were stained *en bloc* in 1% aqueous uranyl acetate overnight at 4°C, then in lead aspartate at 60°C for 30 minutes. After dehydration through a graded ethanol series, samples were infiltrated with acetone and embedded in Epon812 resin (14120, EMS, USA). Ultrathin sections (80 nm) were cut on a Leica EM UC7 ultramicrotome and collected onto 200-mesh nickel grids. For synaptic vesicle quantification in layer 2/3, images were acquired at 14,500X magnification using a Tecnai 20 transmission EM (Thermo Fisher Scientific, USA). Synapses were identified by the presence of clustered presynaptic vesicles, clearly defined synaptic clefts, and an electron-dense PSD.

To enhance visualization of the PSD for excitatory synapse quantification, tissue blocks were processed using a conventional transmission EM protocol. Briefly, blocks were washed in 0.15 M cacodylate buffer, post-fixed in 2% osmium tetroxide, and then stained *en bloc* in 1% uranyl acetate overnight at 4°C. Following serial alcohol dehydration, acetone infiltration, and Epon 812 embedding, ultrathin sections mounted on grids were post-stained with UranyLess (22409, EMS, USA) and lead citrate (22410, EMS, USA). EM images were obtained at 2,500X magnification, and the density of excitatory synapses was estimated using a systematic-random sampling scheme with a 5 μm × 5 μm unbiased counting frame. Random fields within dmPFC layer 2/3 were analyzed in Fiji/ImageJ. Presynaptic bouton area and PSD length were traced manually. Vesicle diameter was measured by outlining the inner membrane of randomly selected vesicles. For vesicle distribution, distances from the center of each vesicle to the presynaptic active zone were calculated by projecting a perpendicular line from the active zone, and vesicles within 400 nm of the AZ were included for analysis. For each synapse, vesicle counts in each distance bin were normalized to the total vesicle number of that synapse, yielding a probability distribution that reflects the relative spatial arrangement of vesicles independent of differences in absolute vesicle number across synapses. The CDF for each synapse was then obtained by computing the cumulative sum across increasing distance bins.

### *in vivo* electrophysiology

#### Data acquisition and spike sorting

For *in vivo* electrophysiological recordings, mice were anesthetized with isoflurane (4% for induction, 0.5–1% for maintenance) and placed in a stereotaxic frame (WPI, USA) with body temperature maintained at 37 °C. For all surgical preparations, the scalp was incised and small craniotomies were made above the target sites for recording or stimulation: dmPFC, AP +1.5 to +1.6 mm from bregma, ML ±0.25 mm from midline, DV −1.1 to −1.3 mm from the skull surface; MD, AP −1.0 mm from bregma, ML ±0.40 mm from midline, DV −3.1 to −3.5 mm from the skull surface. To record dmPFC responses evoked by MD stimulation, 16-channel silicon probes (A1×16-5mm-100-177; NeuroNexus, USA) were used. The probe was slowly lowered until the tip reached a depth of ∼1.5 mm below the cortical surface. Electrical stimulation was delivered as 0.2 ms pulses through an isolated voltage stimulator (DS2A-Mk.II, Digitimer, UK) at intensities of 8–15 V. A small craniotomy was made over the contralateral cortex to implant a microscrew, which served as the reference electrode. During recordings, the brain surface was kept moist by dropwise application of saline.

For recordings during the Y-maze spontaneous alternation task, mice were anesthetized with ketamine (100 mg/kg, i.p.) and placed in a stereotaxic frame (WPI, USA). A silicon probe (A1×16-3mm-100-177-Z16; NeuroNexus, USA) was chronically implanted into the dmPFC for neural recordings during task performance. The probe was slowly lowered until the tip reached a depth of ∼1.5 mm below the cortical surface and was fixed to the skull using dental cement (Super-Bond, Sun Medical, Japan). Ketoprofen (5 mg/kg, subcutaneous) was administered once daily for two consecutive days for postoperative analgesia. Mice were allowed to recover for at least 3–5 days before behavioral testing.

Neuronal activity was amplified and digitized using a PZ5 NeuroDigitizer and RZ5D BioAmp Processor (TDT, USA) at a sampling rate of 24,414 Hz. Signals were band-pass filtered between 0.3 and 5 kHz for single-unit analysis. Spike detection and sorting were performed using Offline Sorter software (Plexon Inc., USA) as described previously^113^. In brief, spike detection was performed by applying a threshold set at four times the baseline noise level, followed by removal of non-physiological artifacts and extraction of a 3.031 ms (74 samples), which included 1.024 ms before the peak time point. The spikes were then aligned to minimize global errors between waveforms. To distinguish units from multiple sources of neurons, we used the automatic scanning of valley seeking algorithm in the principal components space. The algorithm iteratively scans through a range of sorting parameters of valley seeking algorithm, to find the optimal clustering. Outliers were removed from clusters using the software defined quality metrics of Offline Sorter. Units were classified as single units only if they showed significant cluster separation (MANOVA, *p* < 0.05). To further verify the quality of spike sorting, autocorrelograms were generated with 0.1 ms bins using NeuroExplorer (NEX Technology, USA). These autocorrelograms reveal the temporal structure and firing patterns of neuronal activity, which we used to identify refractory periods and assess the isolation of single-unit activity, ensuring the quality and reliability of the spike sorting process. Peri-event firing rates were normalized using Z-score based on each neuron’s overall activity.

#### Single-neuron selectivity analysis

To quantify directional selectivity during the Y-maze task, auROC was computed for each unit by comparing firing rates between left-and right-turn trials within 2 s after delay-zone entry. A selectivity index (SI) was then derived from the auROC value to represent the strength of directional selectivity, independent of turn direction as follows:

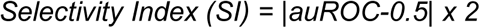

Data acquisition and analysis were performed using TDT processors and Opensorter (TDT, USA), Synapse Software (Synapse Software, USA), and NeuroExplorer (NEX Technology, USA). MATLAB (MathWorks, USA) and GraphPad Prism 6.0 (GraphPad Software, USA) were used for further statistical analyses. All data are presented as mean ± SEM, unless otherwise indicated. Cumulative distributions of SI values were compared between groups using the Kolmogorov–Smirnov test, and one-way ANOVA followed by Bonferroni’s multiple-comparisons test was applied where appropriate. Statistical significance was defined as *p*-values < 0.05, < 0.01, < 0.001 and < 0.0001, as indicated.

#### Population activity analysis

To compare decoding performance across experimental conditions (pre-rKet, rKet, and rKet+C21), recording data were preprocessed as follows. Pseudo-populations were constructed by pooling neurons recorded across sessions. For each condition, the minimum number of simultaneously recorded neurons among sessions was used as the target count. From each session, neurons were randomly selected without replacement until this target number was reached. For decoding, 100 (Fig. 6F) or 20 (Fig. 6H) pseudo-trials were generated per condition by sampling one trial from each neuron’s activity distribution (with replacement across trials). Each pseudo-trial therefore contained one activity value per neuron, preserving statistical independence across neurons. This pseudo-population generation procedure was repeated 1,000 (Fig. 6F) or 100 (Fig. 6H) times to obtain stable estimates of decoding accuracy.

An LDA classifier was trained to determine whether pseudo-population activity could predict the experimental group (pre-rKet, rKet, or rKet+C21). For each iteration, one pseudo-trial served as the test sample and all remaining trials as the training set (leave-one-out cross-validation). This process was repeated so that every trial served once as the test sample. Classification accuracy was defined as the proportion of correctly predicted trials, and the final decoding performance for each condition represented the mean accuracy across 100 pseudo-population iterations.

To determine whether ensemble activity could discriminate between the Delay and Fork epochs, population-level decoding was performed using LDA. To ensure unbiased comparisons across experimental groups and sessions, 100 independent pseudo-populations were generated per group through a two-step sampling procedure:

1. Trial sampling: For each neuron, trials were randomly resampled with replacement (bootstrapping) to equalize trial numbers.
2. Neuron sampling: From the combined pool of all neurons in each group, a fixed number of neurons (matching the smallest count among groups) was randomly selected without replacement to form each pseudo-population.

For each pseudo-population, an LDA classifier was trained to distinguish Delay from Fork activity using repeated random sub-sampling cross-validation (10 repetitions; 50% training, 50% testing). To mitigate potential overfitting, a regularized LDA was used.

Classifier performance was quantified as auROC (Fig. 6F) or SNR (Fig. 6H) of the LDA-projected test data, defined as the absolute difference between the mean projected scores of the two classes divided by their summed standard deviation. This procedure yielded 1,000 SNR values per group (100 pseudo-populations × 10 cross-validation repeats), providing robust estimates of population-level discriminability.

## Acknowledgements

The authors thank members of the Rah laboratory for helpful discussions. This work was supported by grants from the National Research Foundation of Korea (NRF; RS-2023-00266120, RS-2022-NR069145, RS-2024-00508718, RS-2024-00415347, RS-2025-25396400, and RS-2025-02305555) and the Intramural Research Program of the Korea Brain Research Institute (25-BR-01-01, 25-BR-05-05, 25-BR-07-01). The authors also acknowledge the use of ChatGPT (OpenAI, USA) for assistance in correcting grammatical errors. All content was subsequently reviewed and edited by the authors, who take full responsibility for the final manuscript.

## Author information

### Contributions

J.S.K., M.K., J.H.C., and J.C.R. conceptualized the study. J.Y. and I.S.C. performed and analyzed all experiments with contributions from T.L.. G.H.K. acquired and analyzed electron micrographs under the supervision of K.J.L. S.K.B., S.W.B., and J.H.C. contributed to data analysis. J.Y., and J.C.R. wrote the manuscript with input from I.S.C., S.W.B., J.S.K., M.K., and J.H.C.

## Results

## Extended Data

**Extended Data Figure 1.**
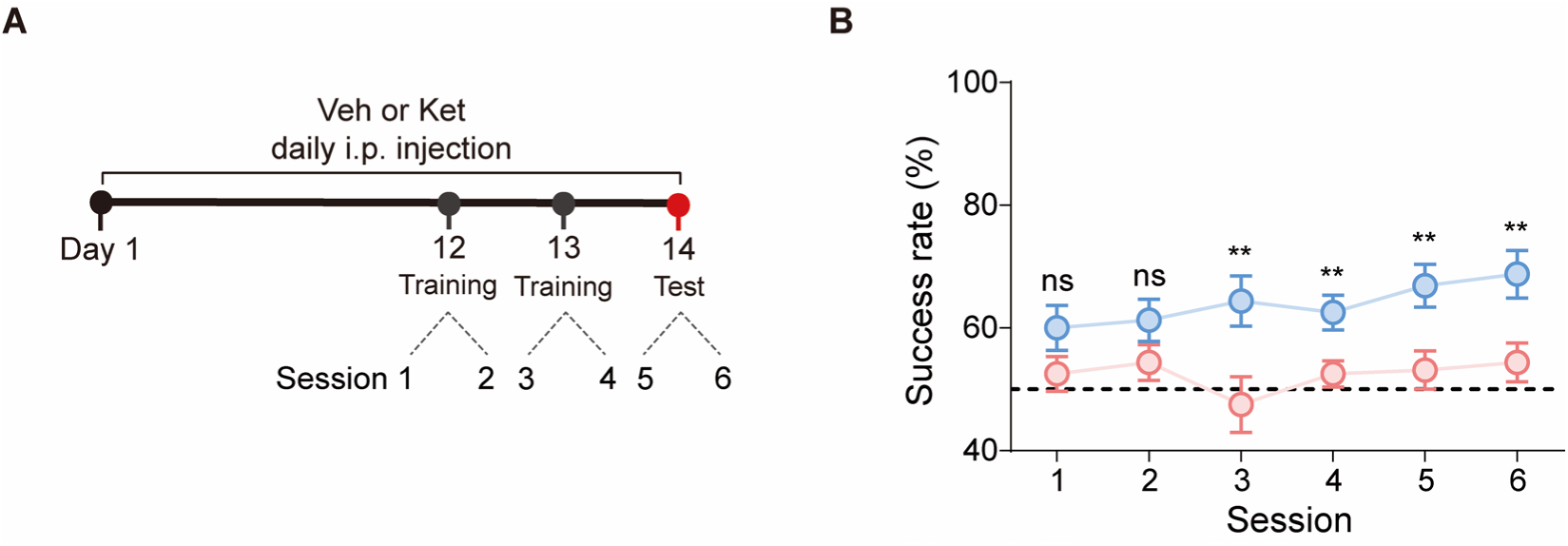
Learning curve across training and testing sessions in the custom-designed Y-maze task. **A**, Experimental timeline showing daily i.p. injections of vehicle or ketamine throughout the 14-day period. Behavioral training was conducted on Days 12–13, and testing on Day 14. Each day consisted of two sessions (Sessions 1–2 on Day 12, Sessions 3–4 on Day 13, Sessions 5–6 on Day 14). Behavioral testing began at least 1 h after injection. **B**, Success rate across six sessions, each comprising 10 trials. Learning performance improved across sessions in control mice but not in rKet mice (Ctrl: 16 mice; rKet: 16 mice). Data are presented as mean ± SEMs; \*\**p* < 0.01 and ns, not significant by unpaired Student’s t-test compared with the Ctrl group.

**Extended Data Figure 2.**
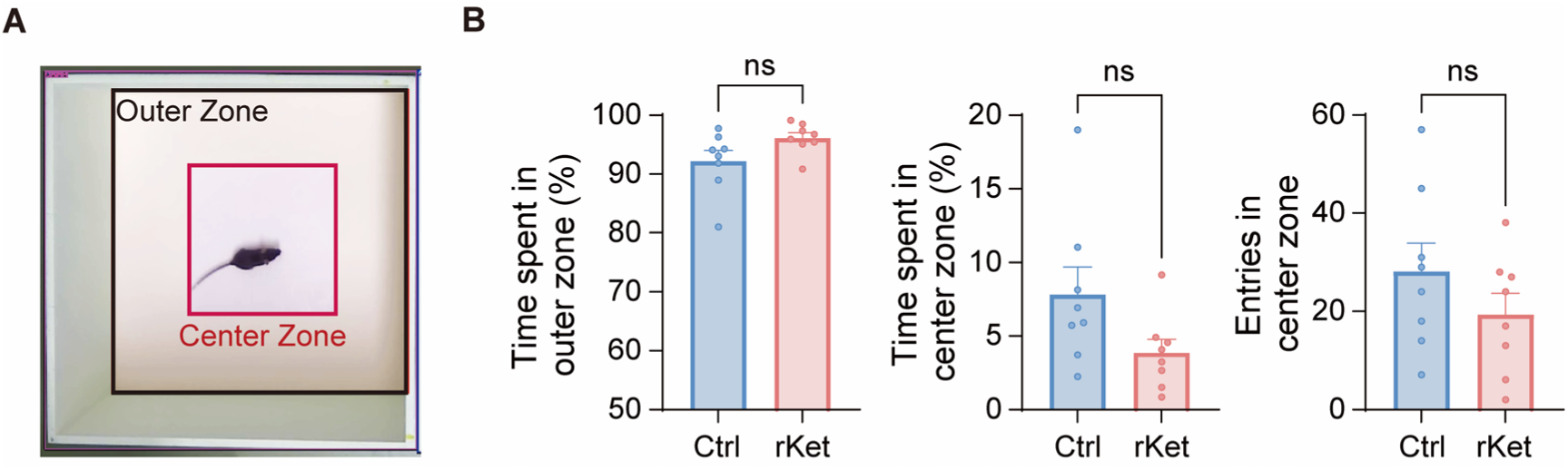
Anxiety-like behavior in the open-field test after repeated ketamine administration. **A,** Schematic of the open field arena showing the center zone (red box) and outer zone (black box). **B,** Quantification of the percentage of time spent in the outer zone (left), center zone (middle), and number of entries into the center zone (right) (Ctrl: 8 mice, rKet: 8 mice). Data is presented as mean ± SEMs; ns, not significant by unpaired Student’s t-test compared with the Ctrl group.

**Extended Data Figure 3.**
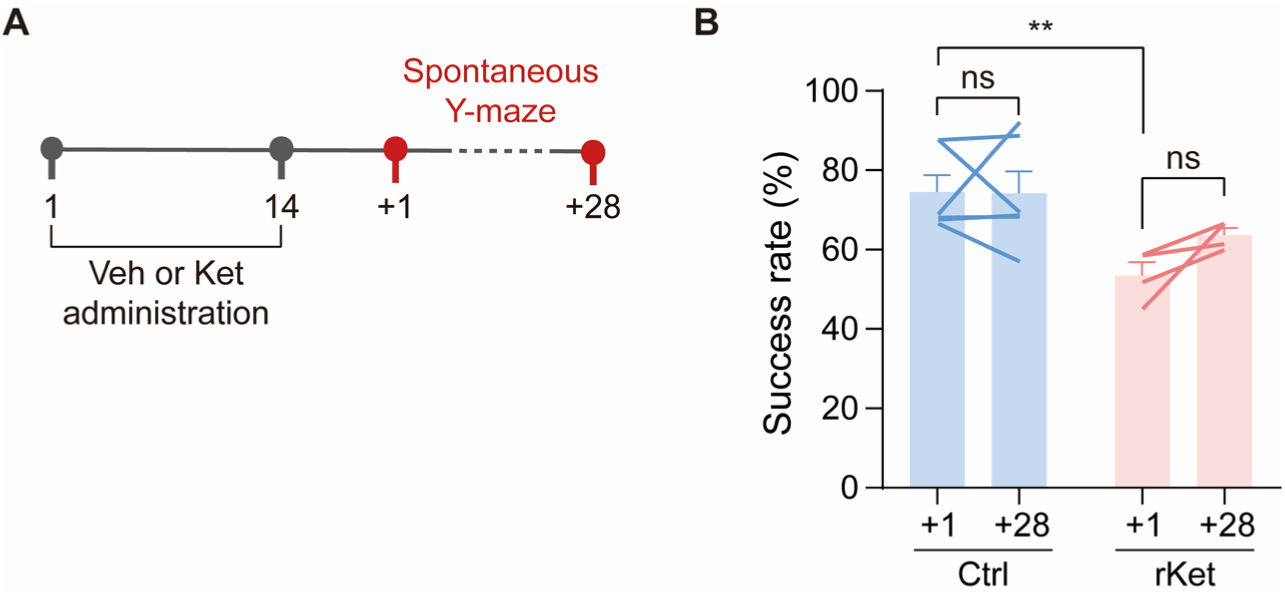
Persistent STM deficit following repeated ketamine administration. **A,** Experimental timeline. Mice received daily vehicle or ketamine injections for 14 consecutive days, followed by a spontaneous Y-maze test on the next day (+1), and an additional test 28 days (+28) later to assess persistence of the deficit. **B,** Success rate in the spontaneous alternation task across the two testing time points (Ctrl: 6 mice; rKet: 4 mice). Data are presented as mean ± SEMs. Within-group comparisons were analyzed with paired Student’s t-tests and between-group comparisons with unpaired Student’s t-tests; \*\**p* < 0.01 and ns, not significant.

**Extended Data Figure 4.**
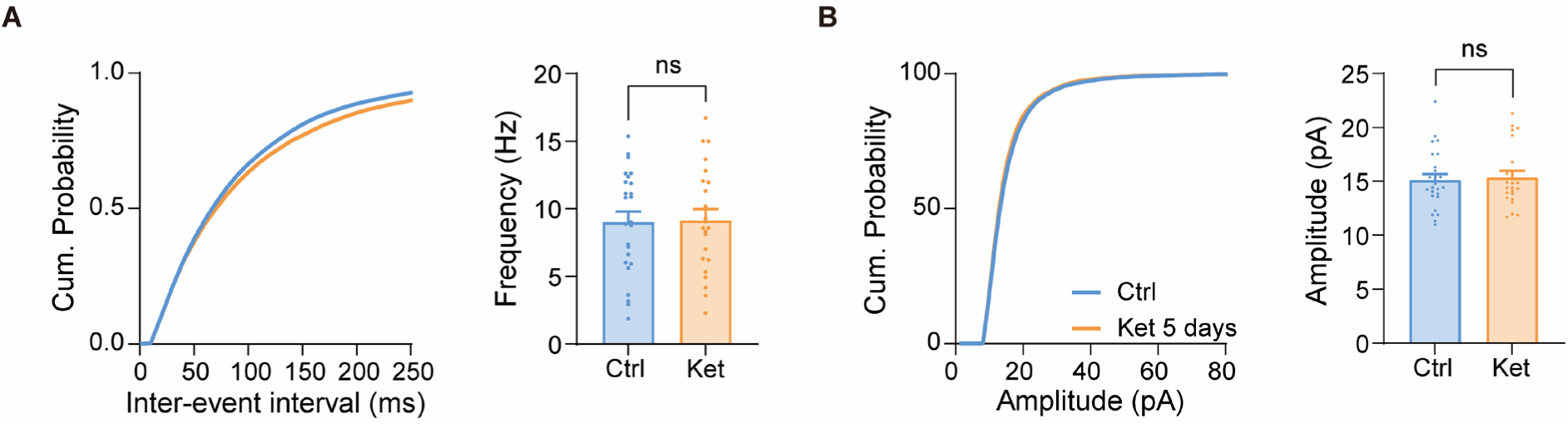
Excitatory synaptic transmission in dmPFC layer 2/3 neurons after sub-chronic ketamine administration. **A,** Cumulative probability plots of inter-event intervals (left) and quantification of average frequency (right) of sEPSCs recorded from dmPFC layer 2/3 neurons after following daily ketamine administration for 5 days. **B,** Cumulative probability plots of sEPSC amplitude (left) and quantification of mean amplitude (right) (Ctrl: 25 cells from 7 mice; Ket: 23 cells from 6 mice). Cumulative distributions were compared using the Kolmogorov–Smirnov test (ns, not significant and \*\*\*\**p* < 0.0001). Data are presented as mean ± SEMs; ns, not significant by unpaired Student’s t-test compared with the Ctrl group.

**Extended Data Figure 5.**
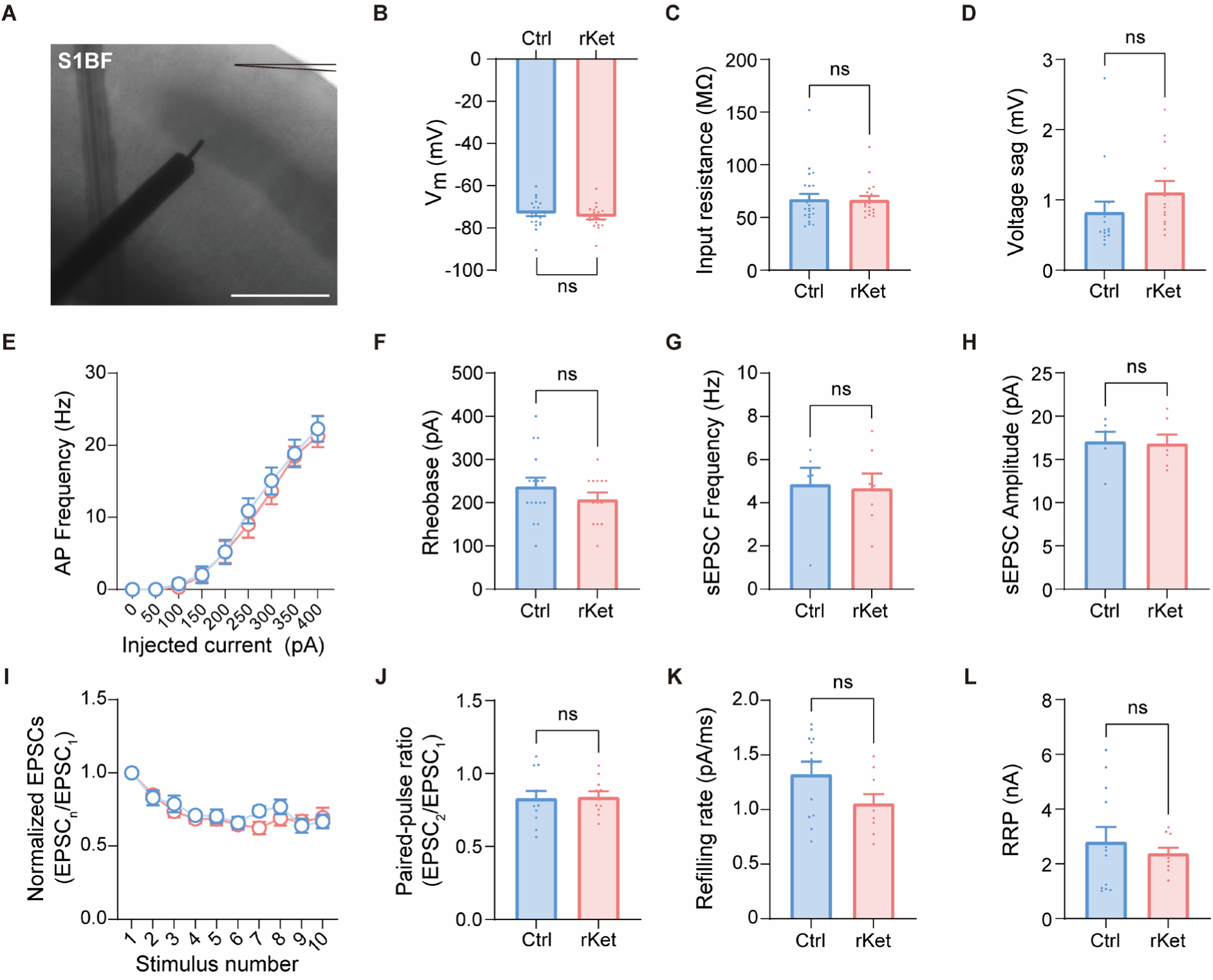
Intrinsic properties and excitatory synaptic transmission in layer 2/3 pyramidal neurons of the primary somatosensory barrel field (S1BF) after repeated ketamine administration. **A,** Representative coronal slice showing a whole-cell recording from a layer 2/3 pyramidal neuron in S1BF. For stimulation experiments, electrical stimuli were delivered to layer 5. Scale bar: 500 µm. **B–F**, Intrinsic excitability of layer 2/3 pyramidal neurons in S1BF following repeated ketamine administration (Ctrl: 24 cells from 6 mice; rKet: 20 cells from 4 mice). **B,** Resting membrane potential (Vₘ). **C,** Input resistance. **D,** Voltage sag. **E,** Action potential (AP) frequency in response to depolarizing current injections (0–400 pA, 50 pA steps, 600 ms). **F,** Rheobase. **G–H**, Spontaneous excitatory synaptic activity recorded from layer 2/3 neurons (Ctrl: 6 cells from 4 mice; rKet: 7 cells from 2 mice). **G**, Mean sEPSC frequency. **H**, Mean sEPSC amplitude. **I–J**, Evoked synaptic responses during 10 Hz stimulation (Ctrl: 12 cells from 6 mice; rKet: 11 cells from 4 mice). **I**, Normalized EPSC amplitudes during 10 Hz stimulation (first to fifth response). **J**, Paired-pulse ratio of evoked EPSCs at a 500 ms interstimulus interval. **K–L**, Synaptic release properties estimated from 50 Hz, 100-stimulus trains (Ctrl: 12 cells from 6 mice; rKet: 9 cells from 4 mice). **K**, Vesicle refilling rate. **L**, RRP size. Data are presented as mean ± SEMs; ns, not significant by unpaired Student’s t-test compared with Ctrl.

**Extended Data Figure 6.**
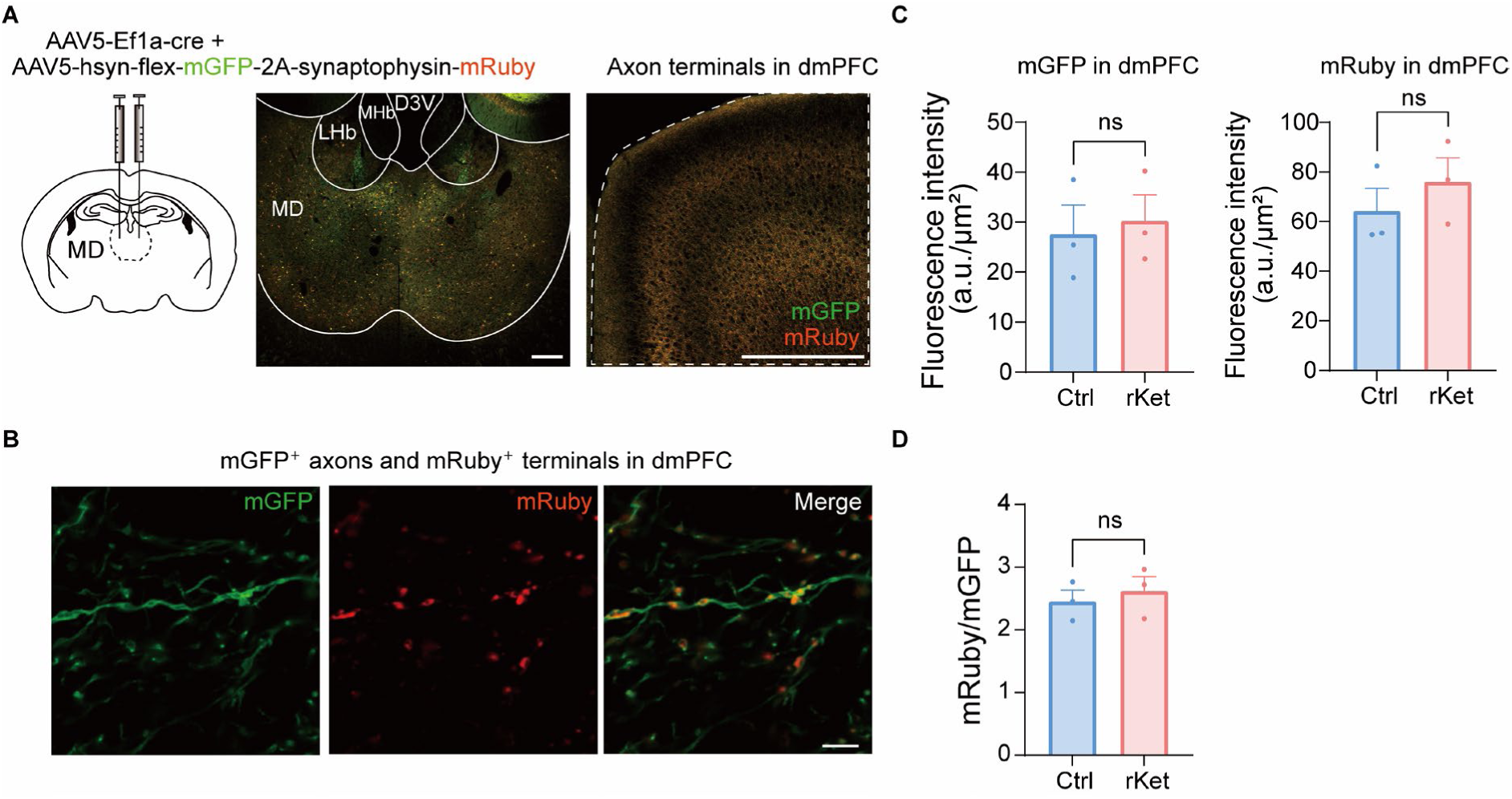
Axonal and synaptic labeling of MD → dmPFC projections after repeated ketamine administration. **A,** Schematic of viral injection strategy. AAV5-hSyn-FLEX-mGFP-2A-synaptophysin-mRuby was bilaterally injected into the MD, and axon terminals were visualized in the dmPFC. Scale bars: left, 200 µm; right, 500 µm. Dotted lines indicate regions of interest (ROIs) used for ImageJ/Fiji analysis. **B,** Higher-magnification images showing axons from dmPFC (mGFP) and their boutons (mRuby) at higher magnification (scale bar: 10 μm). **C,** Quantification of mGFP and mRuby fluorescence intensity (integrated density per area). **D**, Ratio of mRuby to mGFP fluorescence, representing relative synapse-to-projection density. For each mouse, six sections (three per hemisphere) were averaged; n = 3 mice per group. Data are presented as mean ± SEMs; ns, not significant by unpaired Student’s t-test compared with Ctrl.

**Extended Data Figure 7.**
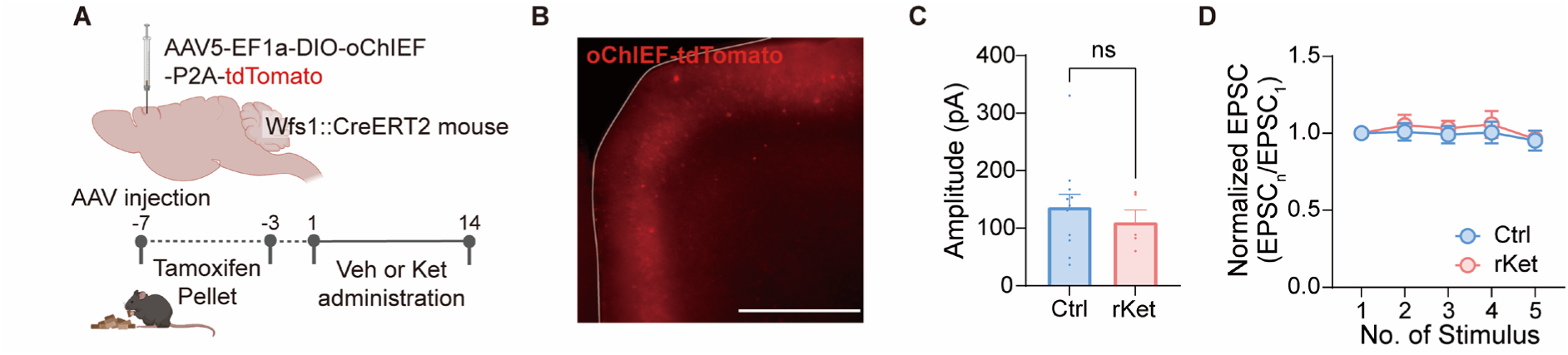
Synaptic release efficacy of cortico-cortical (dmPFC layer 2/3 → layer 2/3) synapses in Wfs1-CreERT2 mice after repeated ketamine administration. **A,** Schematic illustration of experimental design. Cre-dependent oChIEF-tdTomato was injected into the dmPFC of *Wfs1*-CreERT2 mice to selectively express oChIEF in layer 2/3 pyramidal neurons. Tamoxifen pellets were administered for 5 consecutive days to induce Cre activity, followed by repeated vehicle or ketamine administration. **B,** Representative fluorescence image showing oChIEF-tdTomato expression in layer 2/3 of the dmPFC. Scale bar: 500 μm. **C,** Peak amplitude of optogenetically evoked EPSCs recorded at 0.1 Hz stimulation. **D,** Short-term plasticity of evoked EPSCs during 10 Hz stimulation. EPSC amplitudes were normalized to the first response (EPSC_1_) (Ctrl: 12 cells from 4 mice; rKet: 6 cells from 2 mice). Data are presented as mean ± SEMs; ns, not significant by unpaired Student’s t-test compared with the Ctrl group.

**Extended Data Figure 8.**
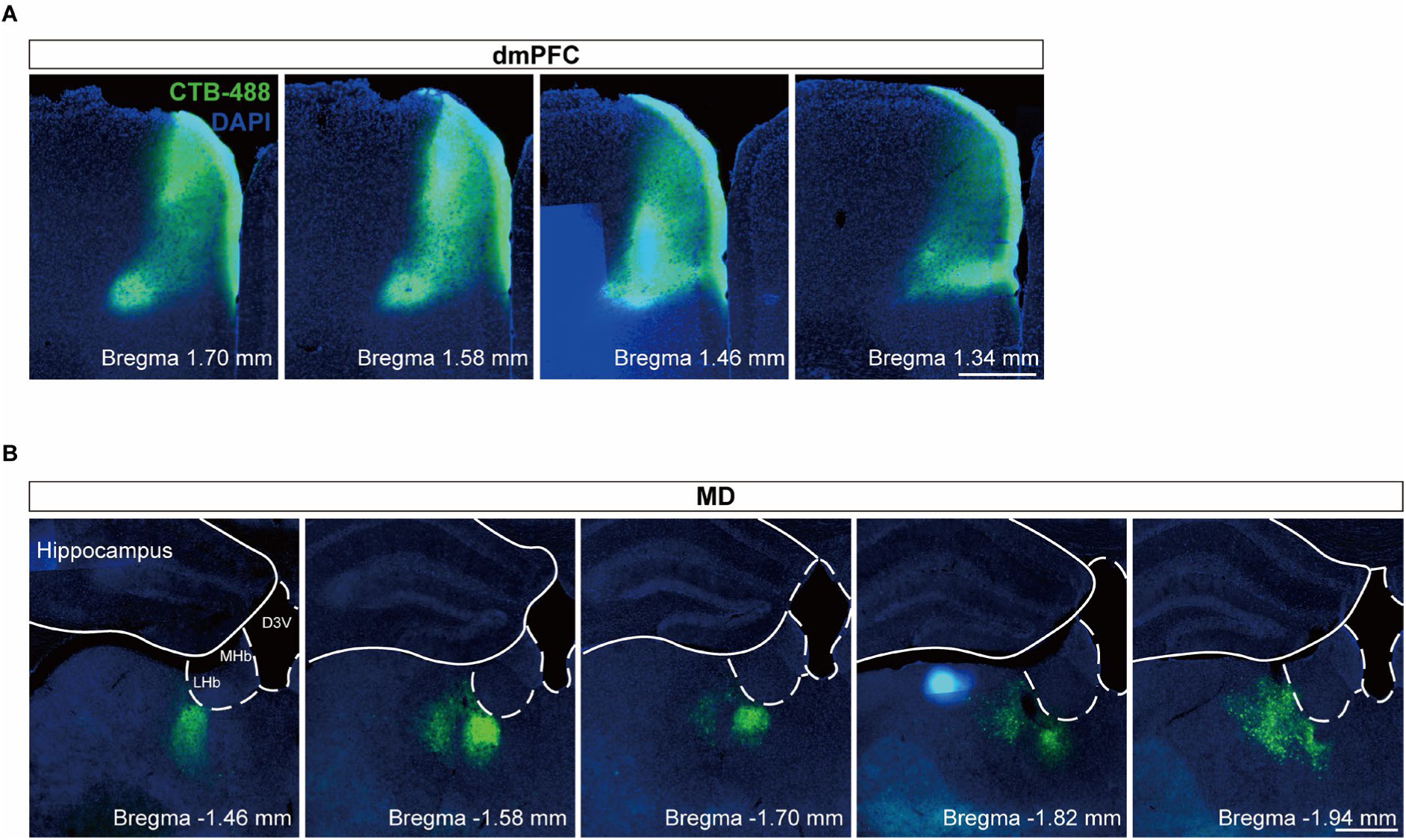
Retrograde tracing reveals MD → mPFC projections. **A,** Injection sites of cholera toxin subunit B conjugated with Alexa-488 in the dmPFC at multiple anterior–posterior levels (AP +1.70 to +1.34 mm). **B**, Retrogradely labeled cell bodies in MD at corresponding posterior coordinates (AP −1.46 to −1.94 mm). D3V, dorsal third ventricle; LHb, lateral habenula; MHb, medial habenula. Scale bar: 500 µm.

**Extended Data Figure 9.**
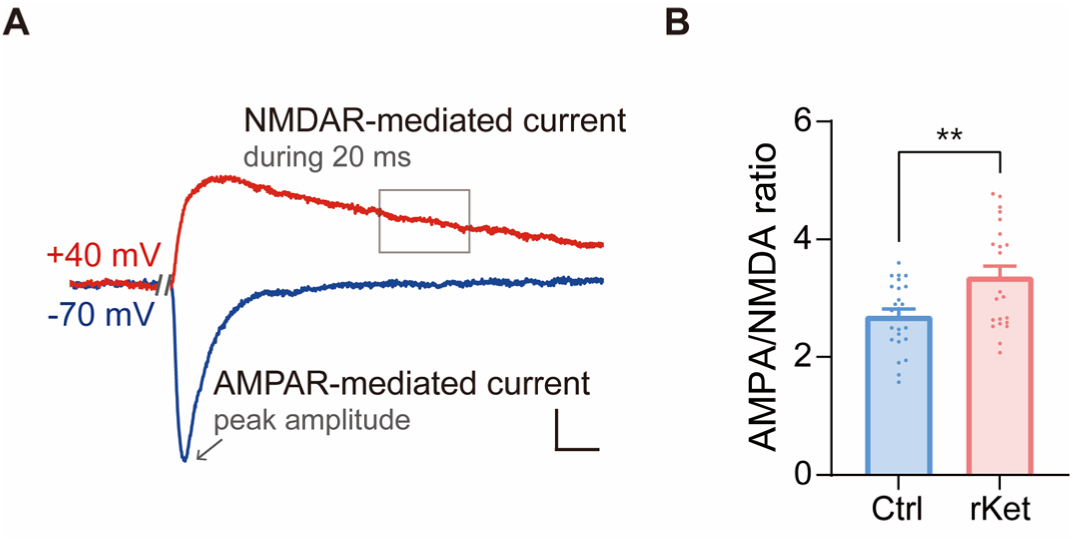
AMPA-NMDA ratio in dmPFC layer 2/3 pyramidal neurons following repeated ketamine administration. **A–B,** AMPA/NMDA ratio measurements in dmPFC layer 2/3 pyramidal neurons. **A,** Representative traces of EPSCs recorded at-70 mV and +40 mV. AMPAR-mediated current was defined as the peak amplitude at −70 mV, and NMDAR-mediated current was quantified as the mean amplitude over a 20 ms window beginning 70 ms after stimulus onset (gray box). Scale bars: 100 ms and 50 pA. **B,** Average AMPA/NMDA ratio (Ctrl: 25 cells from 4 mice; rKet: 22 cells from 4 mice). Data are presented as mean ± SEMs; \*\**p* < 0.01 by unpaired Student’s t-test compared with the Ctrl group.

**Extended Data Figure 10.**
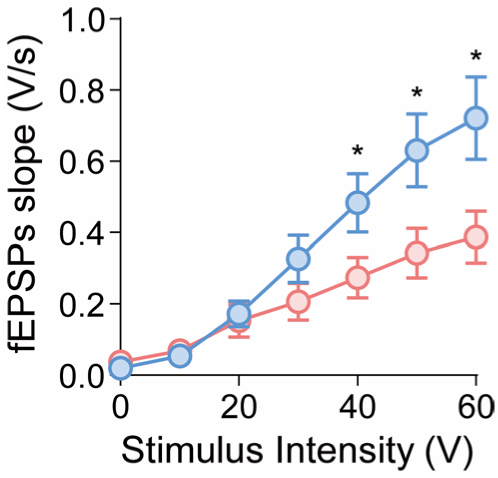
Input–output relationship of dmPFC layer 2/3 synapses following repeated ketamine administration. Field EPSPs (fEPSPs) were recorded from layer 2/3 of the dmPFC while electrical stimulus was delivered to layer 5. The graph shows the relationship between stimulus intensity and fEPSP slope (Ctrl: 19 slices from 7 mice; rKet: 11 slices from 6 mice). Data are presented as mean ± SEMs; \**p* < 0.05 by unpaired Student’s t-test compared with the Ctrl.

